# RAP80 suppresses the vulnerability of R-loops during DNA double-strand break repair

**DOI:** 10.1101/2021.04.23.440542

**Authors:** Takaaki Yasuhara, Reona Kato, Motohiro Yamauchi, Yuki Uchihara, Lee Zou, Kiyoshi Miyagawa, Atsushi Shibata

## Abstract

R-loops, consisting of ssDNA and DNA-RNA hybrids, are potentially vulnerable unless they are appropriately processed. Recent evidence suggests that R-loops can form in the proximity of DNA double-strand breaks (DSBs) within transcriptionally active regions. Yet, how the vulnerability of R-loops is overcome during DSB repair remains unclear. Here, we identify RAP80 as a factor suppressing the vulnerability of ssDNA in R-loops and chromosome translocations and deletions during DSB repair. Mechanistically, RAP80 prevents unscheduled nucleolytic processing of ssDNA in R-loops by CtIP. This mechanism promotes efficient DSB repair via transcription-associated end-joining dependent on BRCA1, Polθ, and LIG1/3. Thus, RAP80 suppresses the vulnerability of R-loops during DSB repair, thereby precluding genomic abnormalities in a critical component of the genome caused by deleterious R-loop processing.

## Introduction

DNA double-strand breaks (DSBs) are a source of genomic instability, such as deletions, insertions, and chromosome translocations, which eventually lead to various disorders including genetic diseases and cancer. Given that the genome exerts its function by being transcribed into RNAs, actively transcribed regions are an important part of the genome. While protein-coding genes occupy ∼3% of the genome, open chromatin regions are estimated as ∼15% of the genome, suggesting that a large region of the genome have a potential to be constantly transcribed (Djebali et al., 2012). Not surprisingly, many roles of RNA in DSB repair at transcription active regions have been suggested; RNA serves as a repair template (Chakraborty et al., 2016; Keskin et al., 2014), forms a DNA-RNA hybrid to recruit repair factors (d’Adda di Fagagna, 2014), and triggers a special type of repair pathway, such as transcription-associated homologous recombination repair (TA-HRR) (Aymard et al., 2014; Ohle et al., 2016; Yasuhara et al., 2018). These mechanisms contribute to maintenance of the actively transcribed regions and protect the most important regions of the genome from genotoxic stresses. However, TA-HRR in S/G2 cells does not function in G1 cells, raising the question of what pathway is responsible for the repair of this class of DSBs in G1 cells.

Intermediates during DNA repair frequently comprise single-stranded DNA (ssDNA), which are a potential vulnerability promoting genomic instability (Chen and Wold, 2014; Wold, 1997). R-loops, which are composed of ssDNA and DNA-RNA hybrids, can form and undergo processing in the proximity of DNA double-strand breaks (DSBs) within transcriptionally active regions (Cohen et al., 2018; Lu et al., 2018; Ohle et al., 2016; Yasuhara et al., 2018). R-loops are generally considered as a potential vulnerability because of the exposed ssDNA regions (Aguilera and Garcia-Muse, 2012). Indeed, accumulating evidence shows that unscheduled R-loop accumulation triggers genomic instability in many contexts (Wells et al., 2019). Given that many nucleases are activated during DSB repair, it is critical to prevent undesired nucleolytic processing of the potentially vulnerable R-loop structure. Nevertheless, how the R-loop-associated vulnerability is overcome during DSB repair remains unclear.

In this study, we identify RAP80 as a key factor suppressing the vulnerability of ssDNA in R-loops during DSB repair, thereby preventing abnormalities in transcribed regions of the genome. We demonstrate that depletion of RAP80 results in accumulation of DNA-RNA hybrids and CtIP and a loss of ssDNA in R-loops. Furthermore, RAP80 directs the repair of DSBs at transcription active regions in non-cycling cells to transcription-associated end-joining (TA-EJ), which requires BRCA1, Polθ and LIG1/3. Thus, we suggest that suppression of the vulnerability of R-loops by RAP80 and TA-EJ are critical mechanisms that protect actively transcribed regions of the genome.

## Results

### RAP80 is a suppressor of genomic abnormalities within gene regions in cancer

To identify the factors precluding genomic instability in gene regions, we exploited The Cancer Genome Atlas (TCGA) database to screen for genes whose low expression is associated with increased deletion risk within gene regions or gene fusion events (Yoshihara et al., 2015). Of 90 DSB-repair factors tested (Table S1), the top hit from both screenings was UIMC1 (RAP80) (Figure 1A and 1B). We identified RAP80 as a possible suppressor of genomic abnormalities occurring in gene regions of the genome. It has been shown that RAP80, a component of the BRCA-A complex, suppresses DSB end resection during homologous recombination repair (HRR) (Coleman and Greenberg, 2011; Hu et al., 2011). We next sought to determine the features of genes in which deletions occurred in the RAP80 low expression group. We selected a set of genes which were highly cell-cycle regulated (59 genes for G1, 36 genes for S and 111 genes for G2 peak expression) (Santos et al., 2015). The samples with low RAP80 expression had an increased risk of deletions at genes highly expressed in G1, but not those highly expressed in S or G2 (Figure 1C). These data led us to hypothesize that RAP80 functions in transcription associated DSB repair in G1 cells to protect genomic regions that are highly transcribed during G1, distinct from its known functions during HRR in S/G2.

**Figure 1.**
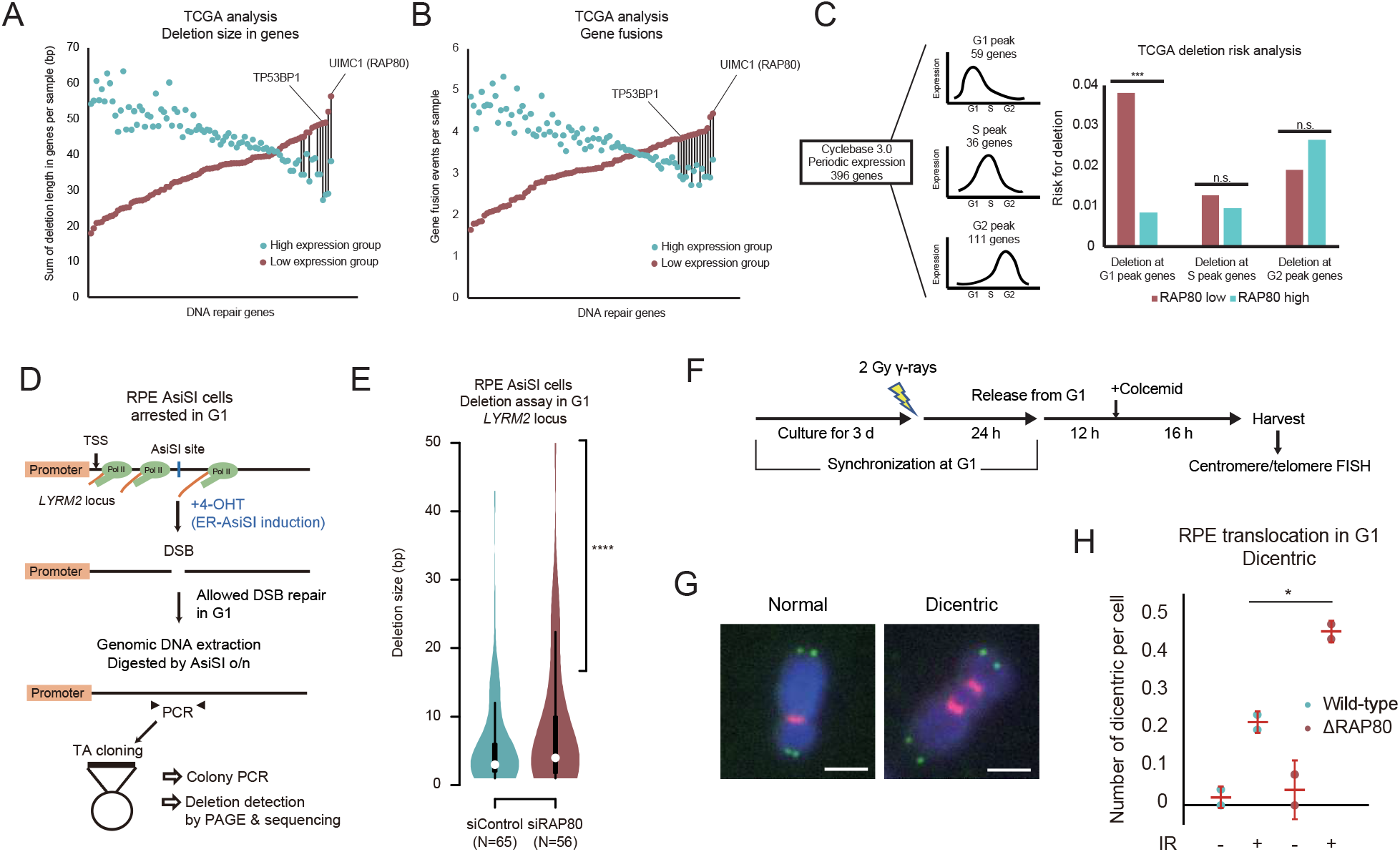
RAP80 suppresses genomic abnormalities at transcribed regions in G1 cells. (A, B) The total number of deleted bases in gene regions of each sample’s genome (A) or the number of gene fusion events per sample (B) was analyzed between the samples with low and high expression of each gene in the TCGA BRCA dataset. Each pair of dots represents low/high expression group of one gene and the black lines between dots indicate significant difference in a two-tailed Welch’s *t*-test. For reference, the dots for TP53BP1 (53BP1) are indicated. (C) The effect of RAP80 levels on the risk of deletions at the G1, S or G2 peak genes was analyzed in the breast cancer samples provided by the TCGA database. Two-tailed *Z*-test. (D) A schematic representation of the deletion analysis at a transcriptionally active locus using G1-synchronized RPE AsiSI cells. (E) The effect of RAP80 depletion on the deletions within the transcriptionally active *LYRM2* locus was analyzed in G1-synchronized RPE AsiSI cells. χ-squared test. For the results of sequencing analysis, see Figure S1A. (F) A schematic representation of the chromosome translocation analysis for G1-synchronized RPE cells. (G) Representative images of a normal chromosome (left) and dicentric chromosome (right) using centromere (red) and telomere (green) fluorescence in situ hybridization (FISH) analysis. Scale bars, 2 μm. (H) The effect of RAP80 depletion on the chromosome translocation frequency was analyzed by counting the number of dicentric chromosomes per cell. Over 2,000 mitotic chromosomes from 50 independent cells were scored per condition in total (mean ± SD, *n*=2 for the data points of two biological replicates).

To consolidate these findings obtained from this database analysis, we examined deletions arising during repair of the AsiSI-induced DSB at the *LYRM2* locus, a transcriptionally active gene, in G1-synchronized RPE cells (Figure 1D). We found that the deletion size at the DSB site is greater in RAP80-depleted cells than wild-type cells (Figure 1E and S1A). These data suggest a role of RAP80 in suppressing deletion size during DSB repair in G1. A similar observation was made in another system detecting deletions in the I-*Sce*I-induced DSB locus in asynchronous H1299 cells (Ogiwara et al., 2011) (Figure S1B and S1C). Next, we examined dicentric chromosomes, which occur as a result of chromosome translocations during DSB repair in G1 (Figure 1F and 1G). RAP80-depleted RPE (ΔRAP80) showed a significant increase in dicentric events induced by ionizing radiation (IR) (Figure 1H), suggesting that RAP80 has a critical role in suppressing translocations during DSB repair in G1. Collectively, these data strongly suggest that RAP80 suppresses genomic abnormalities within transcriptionally active regions in G1 cells.

### Transcription-dependent recruitment of RAP80 to DSB sites

To test how RAP80 is recruited during transcription-associated DSB repair, we employed 730 nm laser irradiation, which preferentially induces DSBs (Reynolds et al., 2013; Yasuhara et al., 2018). We detected recruitment of endogenous RAP80 at early times (∼1 min) after laser irradiation (Figure 2A). Next, we used GFP-tagged RAP80 to follow the kinetics of its recruitment in real time. RAP80 was recruited to laser tracks within 1 min after laser irradiation in G1 cells (Figure 2B). Interestingly, RAP80 recruitment to laser tracks was significantly reduced by treatment with the transcription inhibitor (TRi) (Figure 2C), suggesting that the RAP80 signal in this system represents RAP80 recruitment during transcription-associated DSB repair. Furthermore, RAP80 recruitment was reduced by ubiquitin depletion (Figure 2C). This suggests that RAP80 recruitment to DNA damage sites at early times in G1 cells is dependent on ubiquitin at active transcription sites. Consistent with these findings, a RAP80 mutant which lacks the ubiquitin interacting motif (ΔUIM) was recruited less efficiently to laser tracks compared with full-length RAP80 (Figure 2D). Previous studies suggest that RAP80 is recruited to DNA damage sites dependent on either RNF168-mediated or TRAIP-mediated ubiquitin signaling (Soo Lee et al., 2016; Stewart et al., 2009). However, the RAP80 recruitment at early times detected in our system was not reduced by either RNF168 or TRAIP depletion, suggesting the existence of another ubiquitin signaling pathway to recruit RAP80 (Figure S2A and S2B).

**Figure 2.**
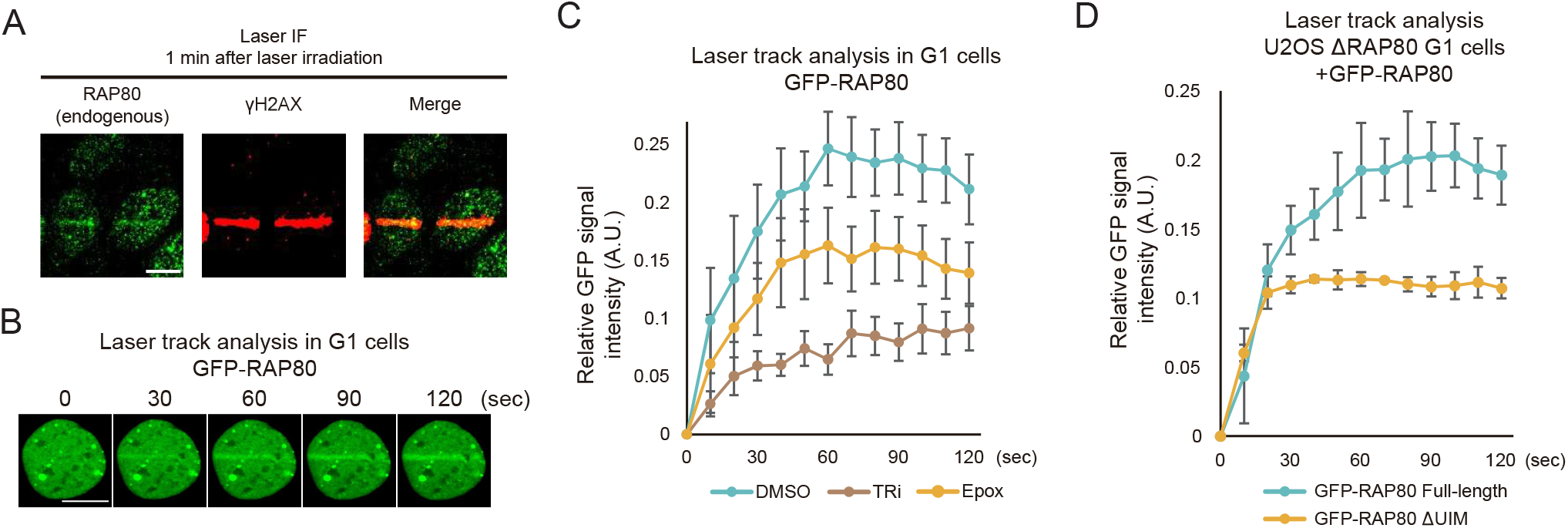
Active transcription-dependent recruitment of RAP80 to DNA damage. (A) The recruitment of endogenous RAP80 to the laser track was analyzed by IF. The cells were fixed 1 min after laser irradiation. Scale bars, 10 μm. (B) Representative images from laser track analysis of GFP-RAP80 recruitment are shown. Scale bars, 10 μm. (C) The effect of TRi (triptolide was used unless otherwise stated), ubiquitin depletion by Epoxomicin (Epox; 10 μM, 30 min) RAP80 recruitment to the laser track was analyzed in G1 U2OS cells (mean ± SEM., *n*=3 for the data points of three biological replicates). (D) The impact of the UIM of RAP80 on recruitment to the laser track was analyzed in G1 U2OS cells (mean ± SEM., *n*=3 for the data points of three biological replicates). To exclude the effect of endogenous RAP80, ΔRAP80 U2OS cells were used in this experiment.

### RAP80 suppresses DNA-RNA hybrid accumulation at DSB sites

Since the recruitment of RAP80 to DNA damage site is dependent on active transcription, it is strongly suggested that RAP80 plays a role in transcription-associated repair of DSBs. We next tracked the accumulation of DNA-RNA hybrids after DSB induction using a GFP-tagged hybrid-binding domain of RNase H1 (GFP-HB) (Bhatia et al., 2014). Interestingly, DNA-RNA hybrids accumulating at the laser-induced DNA damage sites were increased in RAP80-depleted G1 cells (Figure 3A and 3B). In contrast, depletion of MDC1, RNF168, BRCA1 and CtIP, did not affect the accumulation of DNA-RNA hybrids (Figure S3A and S3B). To consolidate these findings, we performed DNA-RNA immunoprecipitation (DRIP). The signals detected by S9.6 antibody DRIP were significantly reduced by RNase H treatment, confirming that these signals are associated with DNA-RNA hybrids (Figure 3C and S3C). The DNA-RNA hybrids accumulating at site-specific DSBs induced by ER-I-*Ppo*I were increased in RAP80-depleted cells at the DSB sites within transcriptionally active genes (Figure 3C). These data suggest a unique role of RAP80 in suppressing accumulation of DNA-RNA hybrids. RAD52 and XPG process DNA-RNA hybrids during DSB repair in S/G2 cells (Yasuhara et al., 2018). Similarly, RAD52 and XPG suppressed accumulation of DNA-RNA hybrids in G1 cells (Figure S3D). Importantly, XPG recruitment to the laser track, but not RAD52 recruitment, was dependent on RAP80 (Figure S3E and S3F). These data indicate that RAP80 coordinates the recruitment of factors involved in processing of DNA-RNA hybrids and reduces their accumulation.

**Figure 3.**
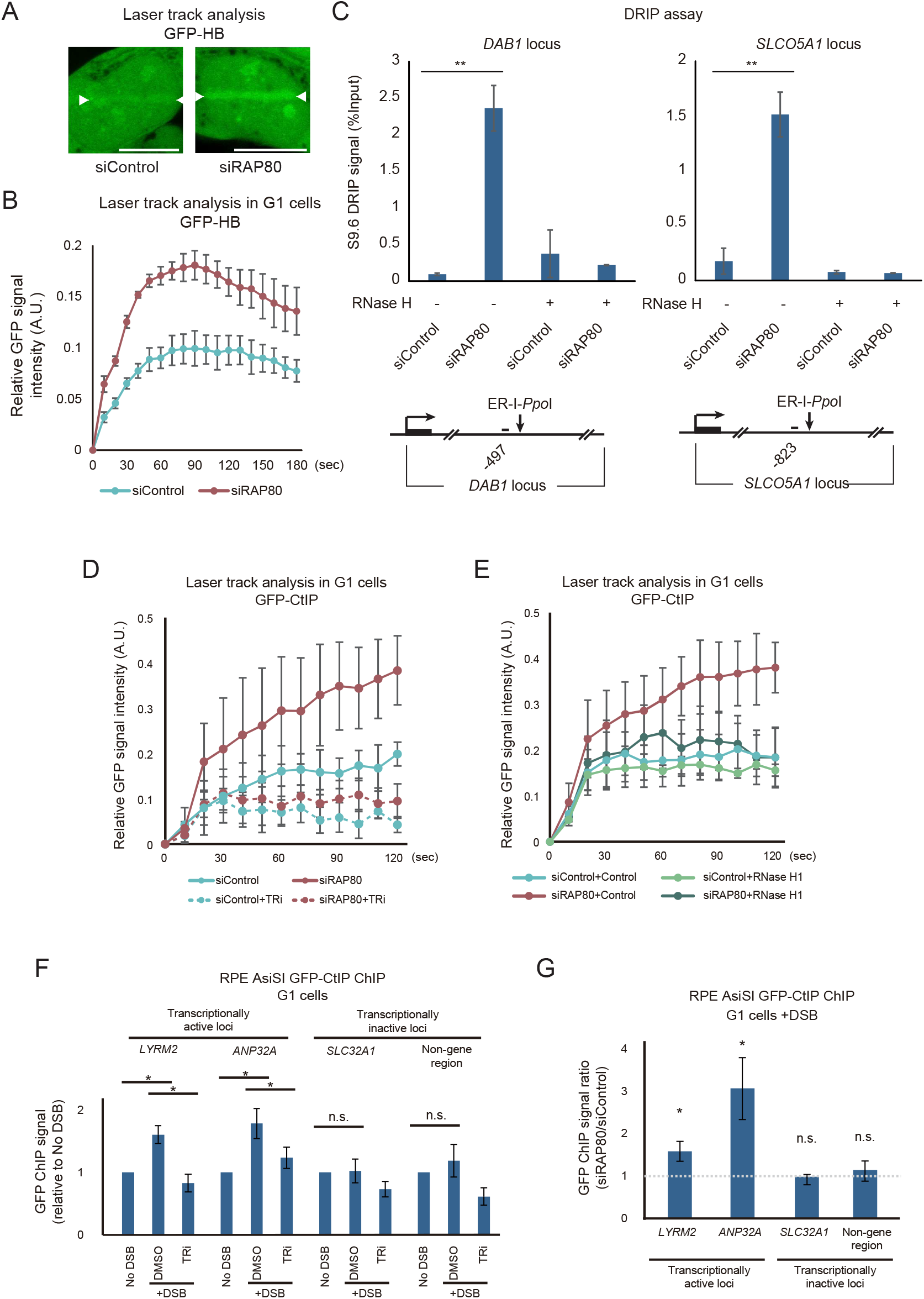
RAP80 suppresses accumulation of DNA-RNA hybrids and CtIP at DSB sites. (A) The accumulation of DNA-RNA hybrids at the laser track 2 min after laser irradiation was analyzed in G1 U2OS cells. White arrowheads, the position of laser irradiation. Scale bars, 10 μm. (B) The effect of RAP80 depletion on DNA-RNA hybrid accumulation at the laser track up to 3 min after laser irradiation was analyzed in G1 U2OS cells by quantifying the GFP-HB signal. Images were recorded with a 10 sec interval (mean ± SEM., *n*=3 for the data points of three biological replicates). (C) The effect of RAP80 depletion on DNA-RNA hybrid accumulation at DSB sites was analyzed by the DRIP assay in HT1080 ER-I-*Ppo*I cells treated with 4-OHT for 1 hour. One representative result of three biological replicates is shown (mean ± SD, *n*=3 for measurements). (D) The effect of RAP80 depletion and TRi treatment on CtIP recruitment to the laser track was analyzed in G1 U2OS cells (mean ± SEM., *n*=3 for the data points of three biological replicates). (E) The effect of exogenous RNase H1 expression in RAP80-depleted cells on CtIP recruitment to the laser track was analyzed as in (D). (F) The recruitment of CtIP to the AsiSI sites after DSB induction and the effect of TRi were compared between transcriptionally active (*LYRM2, ANP32A*) and inactive (*SLC32A1*, non-gene region on chromosome 13) loci using the ChIP assay in G1-synchronized RPE AsiSI cells. The relative signals to the no DSB sample are shown (mean ± SEM., *n*=5-6 for data points from three biological replicates). (G) The effect of RAP80 depletion on CtIP recruitment to DSB sites in G1 cells was analyzed as in (F). The signal ratios (siRAP80/siControl) are plotted (mean ± SEM., *n*=4 for data points of four biological replicates).

### RAP80 suppresses CtIP accumulation at DSB sites

R-loops are a potential source of genomic instability partly because the ssDNA region in their structure can be targeted by DNA nucleases (Aguilera and Garcia-Muse, 2012; Zong et al., 2020). Given that a variety of nucleases function to promote processing of repair intermediates (Lobrich and Jeggo, 2017; Symington, 2014), we hypothesized that accumulation of R-loops upon DSB induction might be vulnerable to nucleolytic incision. A previous study suggested that RAP80 suppresses recruitment of CtIP, a master regulator of DSB end resection, to DSB sites (Coleman and Greenberg, 2011). Interestingly, we detected the recruitment of CtIP to laser-induced DNA damage sites at early times after laser irradiation and this was dependent on active transcription (Figure 3D). Furthermore, RAP80 depletion strikingly increased the amount of CtIP recruited to DNA damage sites in a DNA-RNA hybrid-dependent manner (Figure 3E). A similar increase in DNA-RNA hybrid-dependent recruitment of CtIP was observed in RAD52- and XPG-depleted cells, consistent with DNA-RNA hybrid accumulation in these cells (Figure S3G and S3H). To confirm these findings in a different system, we performed chromatin immunoprecipitation (ChIP) of GFP-CtIP in G1-synchronized RPE AsiSI cells. CtIP was recruited to the DSBs at the transcriptionally active *LYRM2* and *ANP32A* loci, but not to the DSBs at the transcriptionally inactive *SLC32A1* locus or non-gene regions on chromosome 13, and this recruitment was reduced by inhibition of transcription (Figure 3F). Importantly, depletion of RAP80 increased CtIP recruitment to the DSBs at *LYRM2* and *ANP32A* loci, but not the *SLC32A1* locus or non-gene regions (Figure 3G). These data suggest that RAP80 is critical to suppress CtIP accumulation at DSB sites within transcription active regions in G1 cells.

### RAP80 preserves ssDNA in R-loops

The above data prompted us to test whether greater recruitment of CtIP in RAP80-depleted cells promotes unanticipated incision of DSB-induced R-loops and consequent genomic instability. We noticed that XPG recruitment is reduced in RAP80-depleted cells compared with wild-type cells, but this reduction is overridden by additional CtIP depletion (Figure S3F). These data indicate that the double-strand DNA (dsDNA)-ssDNA junction structure at an R-loop, which is normally recognised by XPG (Hohl et al., 2003), is lost in RAP80-depleted cells due to CtIP-mediated incision of the ssDNA within the R-loop, while the DNA-RNA hybrids remain intact. To test this hypothesis, we established a system to track the dynamics of ssDNA in DNA damage-induced R-loops. First, we performed immunofluorescence (IF) analysis of laser irradiated cells using an antibody that recognizes ssDNA or DNA stem loops (Hornick et al., 1998; Ou et al., 2007). The laser tracks observed at very early times (∼1 min) after laser irradiation identified by γH2AX were clearly stained by these antibodies (Figure 4A and 4B), and the staining was reduced by S1 nuclease treatment after fixation, but not RNase A (Figure S4A). Importantly, both antibody signals at the laser tracks were reduced by inhibiting transcription or expressing exogenous RNase H1 (Figure 4C, 4D, S4B, and S4C), suggesting that they represent ssDNA and DNA stem-loop regions of DNA damage-induced R-loops. These data also suggest that at least some, if not all, of the ssDNA region of R-loops can form secondary structures. Interestingly, RAP80 depletion promoted a significant reduction in the amount of ssDNA and DNA stem loops at the DSB site from 1 to 3 min after laser irradiation (Figure 4E). To consolidate these findings on the stability of ssDNA in another system, we stained the laser tracks at early times (∼1 min) after laser irradiation with a BrdU antibody in non-denatured conditions. We found that the laser tracks observed at early times after laser irradiation identified by γH2AX were clearly stained by the BrdU antibody (Figure 4F) and these BrdU signals were significantly reduced by expressing exogenous RNase H1 (Figure 4G). Importantly, RPA, a well-established ssDNA binding complex, was not recruited to laser tracks in G1 cells up to 5 min after laser irradiation, suggesting that the BrdU signals at early times after laser irradiation come from the ssDNA region of R-loops rather than ssDNA caused by DSB end resection (Figure S4D). Strikingly, the ssDNA signals detected by the BrdU antibody were significantly reduced between 1 to 3 min after laser irradiation in RAP80-depleted G1 cells (Figure 4H). Thus, our data strongly support the notion that RAP80 suppresses the loss of ssDNA regions of R-loops during DSB repair in G1 cells.

**Figure 4.**
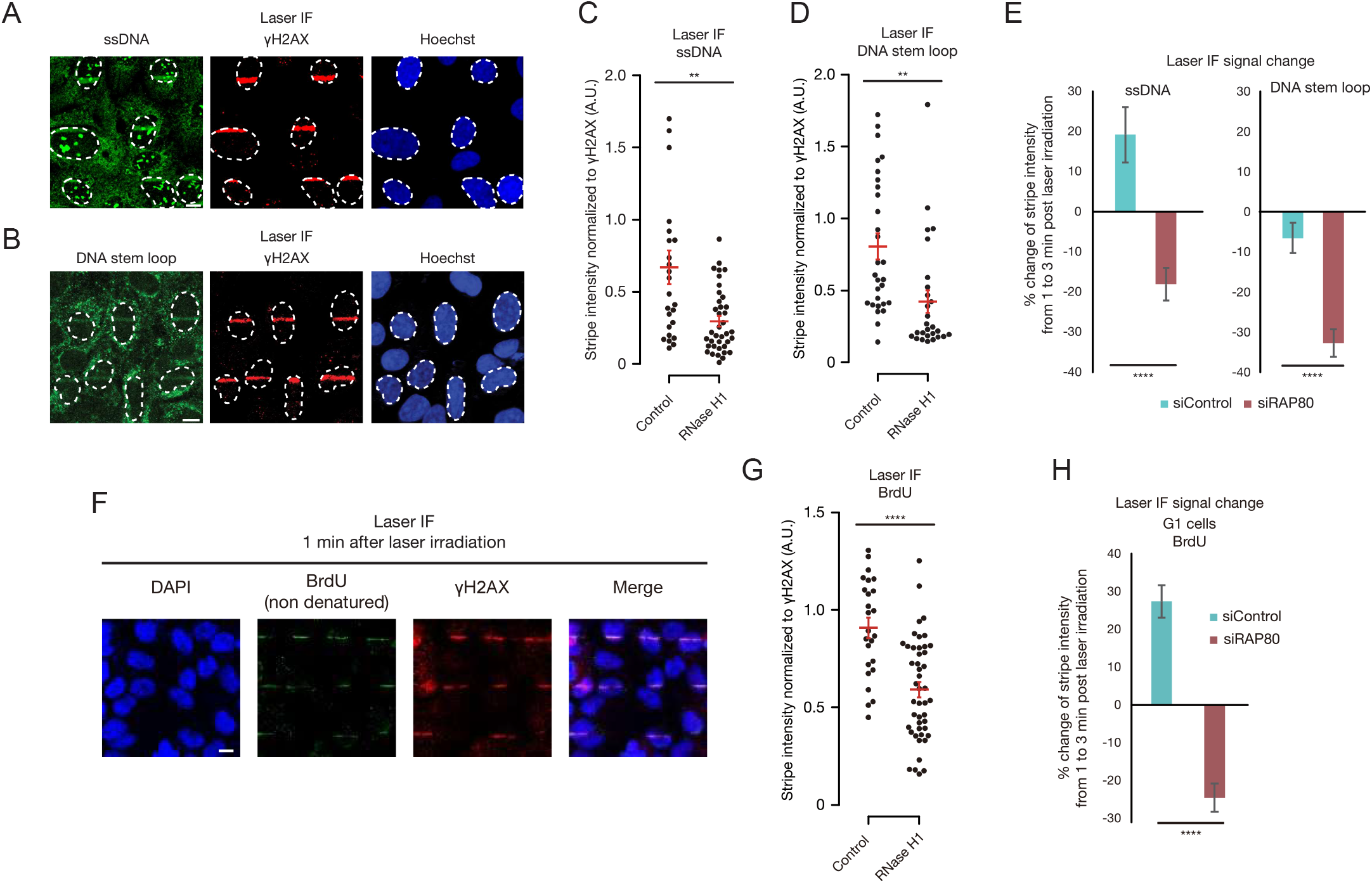
RAP80 preserves secondary-structured ssDNA in R-loops. (A, B) The accumulation of ssDNA (A) or DNA stem loops (B) at the laser track 1 min after laser irradiation was analyzed by IF in U2OS cells. The nucleus of irradiated cells is surrounded by a white dotted line. Scale bars, 10 μm. (C, D) The effect of exogenous RNase H1 expression on ssDNA (C) or DNA stem loop (D) accumulation 1 min after laser irradiation was analyzed by IF (mean ± SEM., *n*=25, 38 (C) and *n*=29, 27 (D), left to right, for data points of independent cells from three biological replicates). (E) The effect of RAP80 depletion on the change in ssDNA or DNA stem loop levels from 1 min to 3 min after laser irradiation was analyzed by IF in U2OS cells (mean ± SEM., *n*=35, 27, 39, 45, left to right, for data points of independent cells from three biological replicates). (F) The accumulation of ssDNA at the laser tracks at 1 min after laser irradiation was analyzed by staining with the BrdU antibody in the non-denatured condition. Scale bars, 10 μm. (G) The effect of exogenous RNase H1 expression on the BrdU signals at the laser tracks was analyzed by IF (mean ± SEM., *n*=25, 46, left to right, for data points of independent cells from three biological replicates). (H) The effect of RAP80 depletion on the change in ssDNA levels from 1 min to 3 min after laser irradiation was analyzed by BrdU IF in mCherry-Geminin-negative G1 cells (mean ± SEM., *n*=50, 45, left to right, for data points of independent cells from three biological replicates).

### RAP80 prevents nucleolytic processing of ssDNA in R-loops by CtIP

Given that CtIP recruitment to DSB sites was increased after RAP80 depletion (Figure 3D-G), we asked whether CtIP is responsible for the loss of ssDNA signals in RAP80-depleted cells. Indeed, the loss of ssDNA and DNA stem loop signals in RAP80-depleted cells were reversed by additional CtIP depletion (Figure 5A). These data strongly indicate that the ssDNA region of DSB-induced R-loops undergoes CtIP-mediated nucleolytic processing in RAP80-depleted cells. To further consolidate these findings, we sought to develop a system to track ssDNA regions in R-loops in real time. A previous report suggested that the ssDNA region at resected DSB ends during HRR forms secondary structures, which are bound by the mismatch repair protein complex MutSβ (MSH2-MSH3 heterodimer) (Burdova et al., 2015). Since DNA stem loops were clearly observed at laser tracks (Figure 4B, 4D, and 4E), we tested whether MutSβ can be used as an indicator of the ssDNA region of DSB-induced R-loops. Interestingly, the dynamics of GFP-tagged MSH2 or MSH3 at laser tracks was fully consistent with the results of the laser IF experiments above; we found that both MSH2 and MSH3 are recruited to laser tracks in G1 cells and reduced by either inhibition of transcription or expression of exogenous RNase H1 (Figure 5B-E). Furthermore, RAP80 depletion significantly reduced the recruitment of MSH2, which was reversed by additional CtIP depletion (Figure 5F), recapitulating the findings in the laser IF experiments above (Figure 4E, 4H, and 5A). Therefore, MutSβ signals at laser tracks likely represent the behavior of ssDNA regions in R-loops. Importantly, neither RAD52 nor XPG depletion decreased MSH2 accumulation (Figure S4E), suggesting that RAD52 and XPG are more likely to process DNA-RNA hybrids rather than ssDNA within R-loops. We also found that CtIP depletion does not affect MSH2 accumulation (Figure S4F), suggesting that CtIP is not involved in ssDNA regulation in the presence of RAP80. Collectively, these data suggest that CtIP is responsible for the loss of ssDNA regions in R-loops in the absence of RAP80.

**Figure 5.**
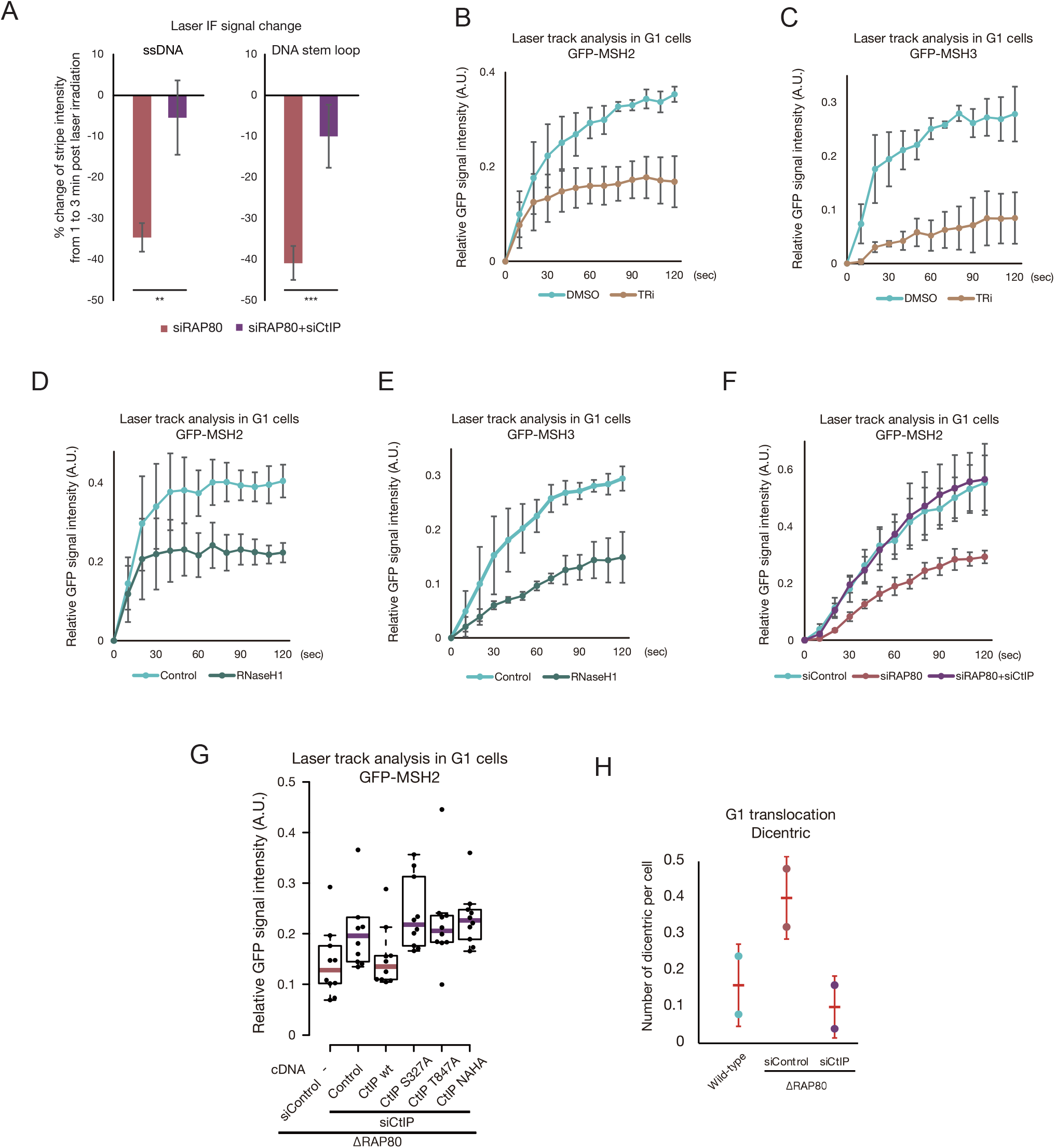
RAP80 suppresses genomic instability caused by CtIP-mediated nucleolytic processing of ssDNA in R-loops. (A) The effect of CtIP depletion in RAP80-depled cells on the change in ssDNA or DNA stem loop levels from 1 min to 3 min after laser irradiation was analyzed by IF in U2OS cells (mean ± SEM., *n*=41, 26, 37, 38, left to right, for data points of independent cells from three biological replicates). (B-E) The effect of TRi (B, C) or exogenous RNase H1 expression (D, E) on GFP-MSH2 (B, D) or GFP-MSH3 (C, E) accumulation at the laser track was analyzed in G1 U2OS cells (mean ± SEM., *n*=3 for the data points of three biological replicates). (F) The effect of CtIP depletion in RAP80-depleted cells on GFP-MSH2 accumulation at the laser track was analyzed in G1 U2OS cells (mean ± SEM., *n*=3 for the data points of three biological replicates). (G) The impact of the CtIP mutants on GFP-MSH2 accumulation at the laser track in RAP80-depleted cells was analyzed at the 1.5 min time point after laser irradiation in U2OS cells and shown as a boxplot (*n*=10 for data points of cells from two biological replicates). (H) The effect of co-depletion of RAP80 and CtIP on the chromosome translocation frequency was analyzed by counting the number of dicentric chromosomes per cell. Over 2,000 mitotic chromosomes from 50 independent cells were scored per condition in total (mean ± SD, *n*=2 for the data points of two biological replicates).

Using this tracking system for the ssDNA regions in R-loops, we next examined whether CtIP catalytic activity is responsible for the incision of ssDNA in RAP80-depleted cells (Makharashvili et al., 2018; Makharashvili et al., 2014). Interestingly, expression of the CtIP endonuclease dead mutant (CtIP NAHA) failed to reduce MSH2 recruitment in RAP80 and CtIP co-depleted cells (Figure 5G, lane 6). Similarly, neither the CtIP S327A mutant, which is defective in interaction with BRCA1, nor the T847A mutant, defective in interaction with the MRE11-RAD50-NBS1 (MRN) complex, reduced MSH2 recruitment in RAP80 and CtIP co-depleted cells (Figure 5G, lanes 4 and 5), suggesting the involvement of CtIP-BRCA1 and CtIP-MRN complexes. Thus, our data demonstrate that CtIP nuclease activity as well as various cofactors stimulated by CtIP are responsible for nucleolytic processing of the ssDNA regions in R-loops in RAP80-depleted cells.

### CtIP-mediated nucleolytic processing of ssDNA in R-loops leads to genomic rearrangements

These data raised the possibility that undesired nucleolytic processing of ssDNA regions in R-loops leads to genomic abnormalities. As shown above, depletion of RAP80 increases chromosome translocation in irradiated G1 cells (Figure 1F-H). We therefore asked whether this increase in chromosome translocation is due to CtIP accumulation and following its nucleolytic processing of R-loops. Strikingly, the additional depletion of CtIP in RAP80-depleted cells reduced dicentric events to the level in control cells (Figure 5H). These data strongly suggest that CtIP-mediated nucleolytic processing of ssDNA regions in R-loops is the cause of genomic rearrangement in RAP80-depleted cells. In turn, RAP80 suppresses CtIP-mediated incision of ssDNA in R-loops, which eventually prevents genomic abnormalities during repair at transcriptionally active regions.

### RAP80 directs DSB repair to TA-EJ

Next, we investigated which DSB repair pathway of G1 cells is regulated by RAP80. First, we examined the efficiency of DSB repair in G1 cells in RAP80-depleted cells. A time course experiment after 1 Gy IR in G1 cells revealed that the number of γH2AX foci at early time points was significantly increased in ΔRAP80 cells (Figure 6A, S5A, and S5B). This repair defect in ΔRAP80 cells at 30 min time point was rescued by expression of full-length, but not ΔUIM, RAP80 (Figure 6B), suggesting that RAP80 recruitment to DSB sites via its UIM is required to promote DSB repair in G1 cells, supporting the model that RAP80 recruitment dependent on its UIM is critical for the DSB response. Consistent with the suggested role of RAP80 in transcription-associated repair, the repair defect in ΔRAP80 cells was reversed by any of three different types of transcription inhibitors (Figure 6C) or exogenous RNase H1 expression (Figure 6D). These data are fully consistent with the notion that RAP80 promotes transcription-associated DSB repair by regulating R-loops. Although additional CtIP depletion in RAP80-depleted cells restores the stability of ssDNA regions in R-loops (Figure 5A and 5G), XPG recruitment (Figure S3F), and dicentric events (Figure 5H), it did not reverse the repair defect at 30 min after IR in ΔRAP80 cells (Figure S5C). These data suggest that the delay in DSB repair in these cells is not completely rescued by additional CtIP depletion (see discussion).

**Figure 6.**
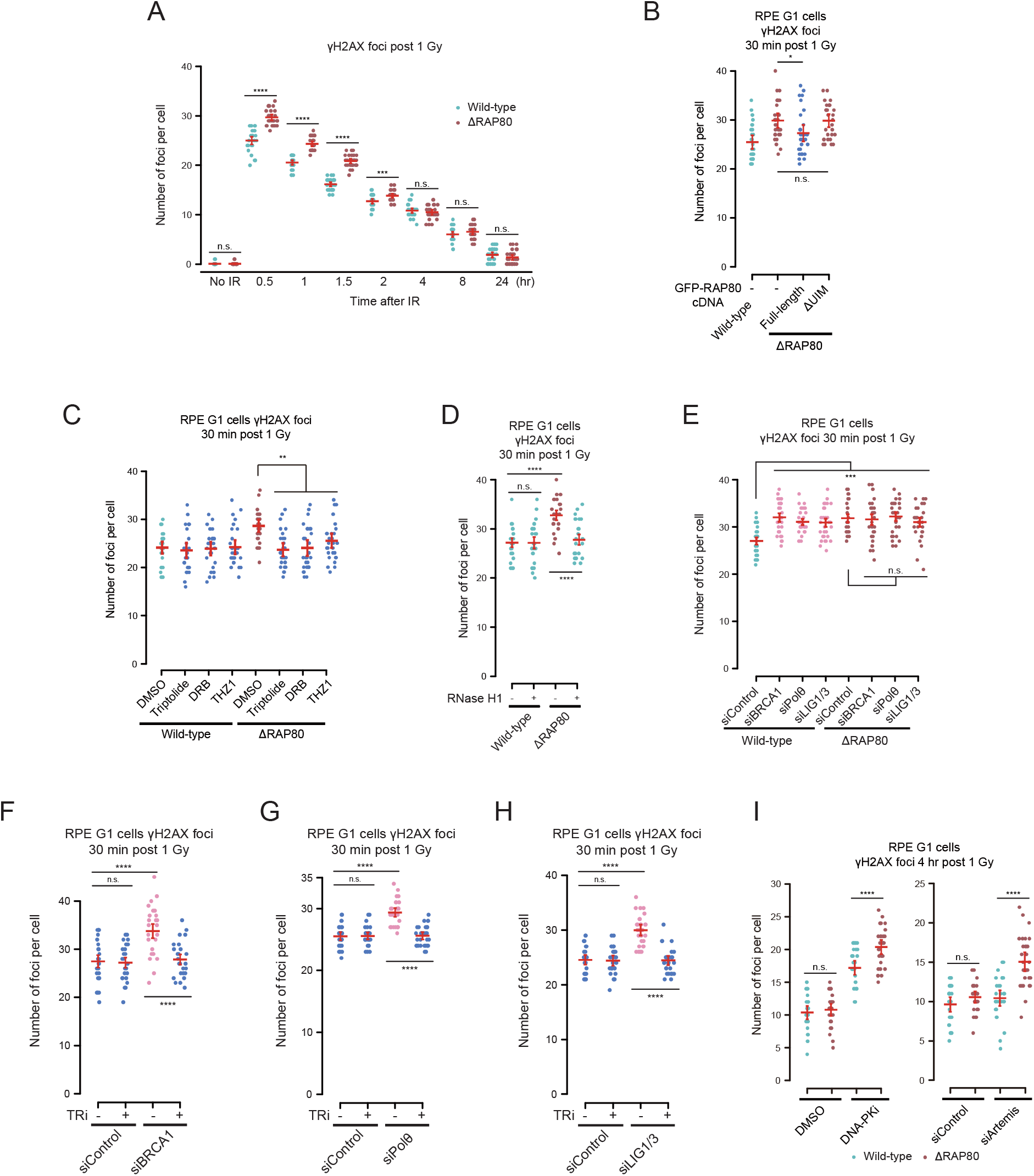
RAP80 coordinates transcription-associated end-joining. (A) The effect of RAP80 depletion on DSB repair kinetics was analyzed by measuring the number of γH2AX foci at the indicated time points in G1 RPE cells. Representative results from three biological replicates are shown (mean with 95% CI, *n*=25 for data points of independent cells). See also Figure S6A. (B) The impact of the UIM of RAP80 on DSB repair efficiency was analyzed by measuring the number of γH2AX foci at 30 min after IR in G1 RPE cells. Representative results from three biological replicates are shown (mean with 95% CI, *n*=25 for data points of independent cells). (C, D) The effect of RAP80 depletion with/without TRi (C) or exogenous RNase H1 expression (D) on DSB repair efficiency was analyzed as in (B). (E) The effect of BRCA1, Polθ or LIG1/3 depletion in wild-type and ΔRAP80 RPE cells on DSB repair efficiency was analyzed as in (B). (F-H) The effect of BRCA1 (F), Polθ (G) or LIG1/3 (H) depletion with/without TRi on DSB repair efficiency was analyzed as in (B). (I) The effect of DNA-PK inhibition (left) or Artemis depletion (right) on DSB repair kinetics in ΔRAP80 RPE cells was analyzed as in (B).

### BRCA1, Polθ, and LIG1/3 are involved in TA-EJ downstream of R-loop processing

We next sought to characterize this RAP80-mediated repair pathway in G1 cells by elucidating factors involved. We identified three major factors BRCA1, Polθ, CtIP, and LIG1/3 involved in transcription-associated DSB repair; depletion of these factors caused a repair defect similar to that caused by RAP80 depletion and was epistatic with RAP80 depletion (Figure 6E-H, S5D, S5E, S6A, and S6B). Dependency on LIG1/3 suggests that the DSBs at transcriptionally active regions are repaired by an end-joining process distinct from canonical non-homologous end-joining. Therefore, we designated this non-canonical pathway as transcription-associated end-joining (TA-EJ). Notably, CtIP also functions in TA-EJ; the wild-type and S327A mutant, but not the T847A and NAHA CtIP mutant, could reverse the repair defect at 30 min after IR in CtIP-depleted cells (Figure S6A and S6B). These data suggest that, although CtIP-mediated incision observed in RAP80-depleted cells is detrimental, CtIP endonuclease activity positively contributes to efficient TA-EJ in wild-type cells. This positive function of CtIP in TA-EJ is similar to what has previously been suggested for CtIP function in DSB repair in G1 cells (Biehs et al., 2017), but is likely unique in that the TA-EJ pathway is initiated by active transcription and R-loops. Nevertheless, CtIP depletion alone did not increase IR-induced dicentric events during G1 DSB repair (Figure S6C) in contrast to RAP80 depletion (Figure 1H), highlighting the importance of RAP80-mediated R-loop processing in suppressing genomic abnormalities.

Since BRCA1 depletion did not lead to the accumulation of DNA-RNA hybrids (Figure S3B), we asked whether BRCA1 is required downstream of R-loop processing by RAP80, RAD52 and XPG. In RAD52- and XPG-depleted cells, the repair defect at 30 min after IR was not obvious (Figure S6D, lanes 1, 4, 7), suggesting that a backup repair pathway quickly rejoins the DSBs in these cells (see below). RAP80 depletion delayed DSB repair in wild-type, RAD52- and XPG-depleted cells (Figure S6D, lanes 2, 5, 8) probably because RAP80 is the most upstream regulator within these factors. In contrast, BRCA1 depletion caused repair defects at 30 min after IR in wild-type cells, but not in RAD52- and XPG-depleted cells (Figure S6D, lanes 3, 6, 9). These data suggest that RAP80 is upstream of RAD52 and XPG, whereas BRCA1 is downstream of RAD52 and XPG.

### Defective TA-EJ is backedup by an Artemis-dependent pathway

Importantly, the repair defect in ΔRAP80 cells observed at 30 min after IR was gradually diminished at later time points (Figure 6A), suggesting the presence of a backup repair pathway. Strikingly, inhibition of DNA-PK activity or depletion of Artemis caused persistence of the repair defect in ΔRAP80 cells at 4 hours after IR (Figure 6I), demonstrating that this backup pathway is dependent on DNA-PK and Artemis. In RAD52- and XPG-depleted cells, the repair defects at 30 min after IR were observed only when Artemis is depleted (Figure S6E), suggesting that defective TA-EJ caused by RAD52 or XPG depletion is backedup by an Artemis-dependent pathway. As expected from the functions of RAD52 and XPG in R-loop processing, the repair defects observed in RAD52/XPG and Artemis co-depleted cells were rescued by exogenous RNase H1 expression (Figure S6F), indicating RAD52 and XPG are dispensable in this pathway in the absence of R-loops. Collectively, these data demonstrate that Artemis is commonly used in the backup pathways for TA-EJ.

## Discussion

### R-loop processing is an integral part of DSB response

We dissected the significance of the two strands of R-loops, i.e., the DNA-RNA hybrid and ssDNA, in DSB repair, which are processed individually and distinctly. The protection of the ssDNA region by RAP80 precedes the processing of the DNA-RNA hybrid by RAD52 and XPG. If DSB-induced R-loops are caused by collision between ongoing transcription and the DSB end, the ssDNA region of the R-loop after incision of DNA-RNA hybrids by XPG can serve as a 3’ overhang, which is a primary substrate for Polθ-mediated DSB repair (Figure 7) (Kent et al., 2015; Mateos-Gomez et al., 2017). Given that alternative end-joining pathways generally require CtIP for DSB end resection (Zhang and Jasin, 2011), the roles of CtIP in Polθ-mediated TA-EJ are reasonable. In contrast, in the absence of RAP80, CtIP accumulates at DSB sites and incises ssDNA in R-loops collapsing the dsDNA-ssDNA junction structure, which is normally recognized by XPG, thereby leading to an accumulation of DNA-RNA hybrids. These data suggest that unscheduled R-loop processing by nucleolytic enzymes can result in deleterious consequences for the genome. Although it is still unknown how RAP80 suppresses CtIP accumulation, our data indicate that DNA-RNA hybrids recruit CtIP in RAP80-depleted cells. It is possible that RAP80 keeps R-loops unexposed by recruiting some factors in the proximity of DSBs.

**Figure 7.**
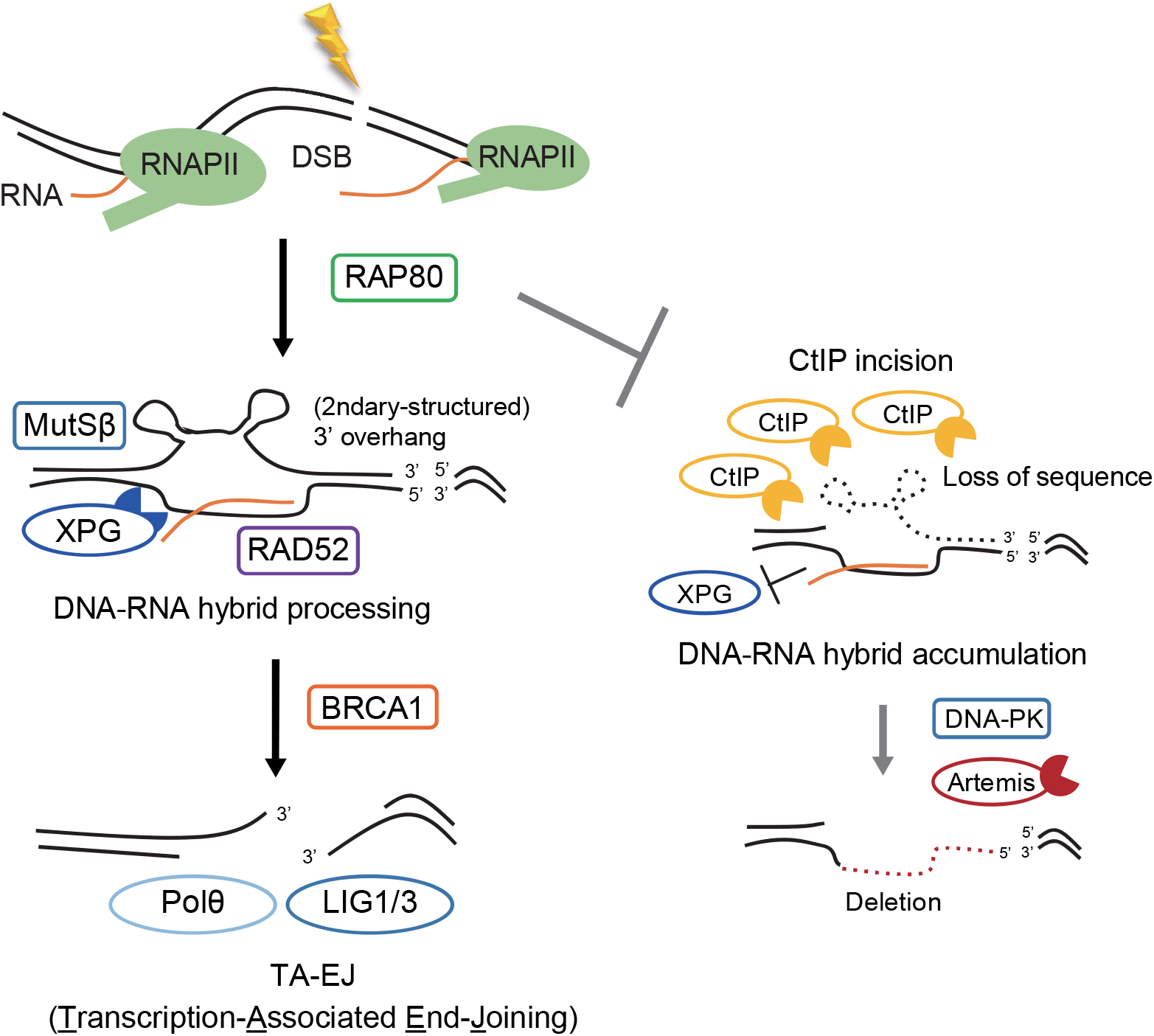
The model for TA-EJ initiated by RAP80. A schematic model for DSB repair in transcriptionally active regions by TA-EJ. See the text for detail.

### TA-EJ minimizes aberrant repair consequences at transcribed regions

Cells must have special DNA repair mechanisms by which they maintain the stability of the most important regions of the genome, i.e., transcribed regions. In this study, our data demonstrate that the interplay between the transcription and DSB repair machinery protects the actively transcribed regions of the genome. The dependency of TA-EJ on RAD52, XPG as well as BRCA1, CtIP, Polθ, LIG1/3 suggests that the ends of DSB within transcribed regions require some extent of resection before end joining. It is therefore possible that the nature of DSB ends at transcribed regions precludes these DSBs from being precisely repaired. Meanwhile, a defect in TA-EJ significantly increases the deletion size and chromosome translocations. Thus, TA-EJ is the mechanism that minimizes the aberrant repair consequences at transcribed regions.

The formation of R-loops at the DSB site and their proper processing triggers at least two pathways: TA-EJ and TA-HRR. Although RAD52 and XPG process DNA-RNA hybrids in DSB-induced R-loops throughout the cell cycle to promote both TA-EJ and TA-HRR, RAP80 appears to function differently; RAP80 promotes TA-EJ and suppresses TA-HRR (Yasuhara et al., 2018). The fact that RAP80 depletion in S/G2 cells increases the fraction of DSBs repaired via TA-HRR may explain why the contribution of RAP80 to genome stability appears mostly in the G1 phase when HRR is inexecutable. We propose that genome stability in transcriptionally active regions is maintained by two distinct, but complementary mechanisms throughout the cell cycle.

## Acknowledgments

We thank Prof. Penny A. Jeggo for critical discussion. The H1299 dA-3 cell line was a generous gift from Dr. Ogiwara. RPE-hTERT cells harbouring the OsTIR1 expression cassette (Venegas et al., 2020) was a generous gift from Dr. Kanemaki. We thank Yoshimi Omi, Akiko Shibata, Yoko Hayashi, Yukihiko Yoshimatsu, Itaru Sato, Shiho Nakanishi, Naho Takashima, Hiroko Iino, and Yoshihiko Hagiwara for assisting with the lab work and other supports. This work utilized the core research facility of Center for Biology and Integrative Medicine, The University of Tokyo, which was organized by The University of Tokyo Center for NanoBio Integration entrusted by Ministry of Education, Culture, Sports, Science and Technology (MEXT) Japan. This work was carried out under the support of Isotope Science Center, The University of Tokyo. A part of this study was conducted through the Joint Usage/Research Center Program of the Radiation Biology Center, Kyoto University. Radiation Biology Center is a joint usage research center certified by MEXT Japan. AS is a visiting associate professor of Radiation Biology Center, Graduate School of Biostudies, Kyoto University. This work was supported by JSPS KAKENHI Grant Number JP18K18191 to T.Y., JP15H04902 and JP15K14376 to K.M., JP26701005 and JP17H04713 to A.S., the Takeda Science Foundation to T.Y. and A.S., the Uehara Memorial Foundation to A.S., the Mitsubishi Foundation to A.S. and the Yamada Science Foundation to T.Y.

## Author contributions

T.Y., R.K., and A.S. conceived and designed the study discussing with K.M. The experiments were performed by R.K., M.Y., Y.U. and T.Y. T.Y., R.K., M.Y., T.O., and A.S. analyzed the results. T.Y., L.Z., K.M. and A.S. wrote and edited the manuscript. A.S. supervised the study.

## Methods

### Cell lines

All cell lines were cultured in a 37 °C incubator at 5% CO2 and in media supplemented with 10% fetal bovine serum and penicillin/streptomycin. The RPE-hTERT (Clontech) cell line was cultured in DMEM/F-12 medium, U2OS in McCoy’s 5A medium, HT1080 ER-I-*Ppo*I cell line in α-MEM medium and H1299 dA-3 cell line in MEM medium. The U2OS mCherry-Geminin cells, RPE-hTERT ΔRAP80 and ΔRAD52 cells were described previously (Yasuhara et al., 2018). For generation of RPE-hTERT ΔArtemis and U2OS ΔRAP80 cells, the All-in-One CRISPR-Cas9D10A vector (generous gift from Prof. Steve Jackson) (Chiang et al., 2016) with the gRNAs specific for the target gene was transfected and sorted by a FACSAria II cell sorter (Becton Dickinson). The clones were confirmed by western blot. For generation of RPE-hTERT AsiSI cells, an ER-AsiSI-AID vector was transfected into RPE-hTERT cells harboring the OsTIR1 expression cassette (Venegas et al., 2020), and cells stably expressing ER-AsiSI-AID were selected by G418.

### Laser track analysis

For real-time laser track analysis, the U2OS mCherry-Geminin cells were plated on a 35 mm glass bottom dish (Matsunami) and transfected with the indicated plasmids. On the day of analysis, cells were irradiated with the 730 nm laser with approximately 500 nm in width using a LSM510 microscope (Zeiss) after incubated with 10 μg mL^-1^ Hoechst33342 (Wako) for 10 min. For TRi treatment, cells were treated with triptolide (3 μM) for 30 min prior to laser irradiation. Throughout the analysis, cells were kept at 37 °C using a temperature control chamber (Zeiss). In most experiments, GFP-positive and mCherry-negative (G1) cells were analyzed. Images were collected at a 10 sec interval and the GFP signal on the laser track was quantified using a Zen software (Zeiss). At least five cells per condition were examined. In all panels, the mean and standard error of three biological replicates are shown. For laser IF analysis, cells were fixed with either 100% MeOH at -20 °C or 4% PFA at room temperature for 5 min before incubation with the primary antibodies overnight at 4 °C. Images were captured using a LSM 510 META or LSM 880 (Zeiss). The signals on the laser track were quantified using an Image J software and normalized with those of γH2AX.

### Immunoprecipitation

Cells were treated with Zeocin (3 mg mL^-1^) for 1 hour and lysed in IP buffer (50 mM Tris-HCl, 150 mM NaCl, 1 mM EDTA, 0.5% NP-40 supplemented with Halt Protease Inhibitor Cocktail (Thermo Fisher Scientific) and Phosphatase Inhibitor Cocktail 3 (SIGMA)). After brief sonication, lysate was incubated with primary antibody for 2 hours at 4 °C in rotation with or without EtBr (50 μg mL^-1^) before adding Dynabeads Protein A/G for 30 min. The beads were washed 4 times with IP buffer, boiled in SDS sample buffer for 5 min, and analyzed by western blot. For chromatin immunoprecipitation, cells were fixed with 2% paraformaldehyde for 10 min followed by addition of 1 M Glycine (final 125 mM) for 5 min. After lysis of the cells in IP buffer, chromatin was fragmentated by sonication using water bath sonicator at 4 °C. The lysate was incubated with primary antibody for 2 hours at 4 °C in rotation before adding Dynabeads Protein A/G for 30 min. After washing 4 times with IP buffer, chromatin was eluted in 100 mM NaHCO3, 1% SDS for 10 min at room temperature. The protein-DNA crosslink was reversed by incubating 65 °C overnight. The eluate was purified using QIAquick PCR purification kit (Qiagen) and analyzed using LightCycler 480 (Roche).

### DRIP assay

The ER-I-*Ppo*I cells treated with 4-OHT (2.5 μM) for 1 hour were collected for extraction of the total nucleic acids using DNeasy Blood & Tissue Kits (Qiagen) or PureLink(tm) Genomic DNA Mini Kit (Thermo Fisher Scientific) according to the manufacturer’s instructions. The eluted DNA was incubated with or without RNase H (New England Biolabs) at 37 °C overnight. DNA-RNA hybrids were immunoprecipitated from the total nucleic acids using the S9.6 antibody overnight. The immunoprecipitate was purified using Illustra GFX PCR DNA and Gel Band Purification Kits (GE Healthcare) or QIAquick PCR purification kit (Qiagen) and analyzed using a 7500FAST instrument (Applied Biosystems) or LightCycler 480 (Roche).

### Foci analysis

For foci analysis in irradiated RPE G1 cells, cells were grown on a coverslip and used 1 day after 100% confluent. Cells were treated with 5-ethynyl-2’-deoxyuridine (EdU, 10 μM), TRi (triptolide, 3 μM; DRB, 100 μM; THZ1, 1 μM) or DNA-PK inhibitor (NU7026, 10 μM) for 1 hour before IR and subsequent fixation with 4% PFA in PBS and permeabilized with 0.1% SDS, 0.5% Triton-X in PBS at the indicated time point. Cells were incubated with the γH2AX antibody for 1 hour at 37 °C, followed by 1 hour incubation with secondary antibodies. For visualization of incorporated EdU, a click chemistry procedure was performed. For IR, a CellRad (Faxitron) was used.

### Deletion assay

For the deletion assay in G1 RPE AsiSI cells, 4-OHT (300 nM) was added to induce ER-AsiSI-AID dependent DSBs at 96 hours after RAP80 siRNA transfection. Two days after the addition of 4-OHT, genomic DNA was isolated by genomic DNeasy Blood & Tissue Kit (Qiagen). The genomic DNA (1 μg) was digested with 20 U AsiSI at 37 °C overnight and purified with phenol-chloroform extraction followed by ethanol precipitation. The *LYRM2* locus containing an AsiSI cut site was amplified by PrimeSTAR® Max DNA Polymerase (Takara). The PCR product was purified with NucleoSpin Gel and PCR Clean-up (Takara) and digested with AsiSI at 37 °C overnight. The digested PCR product was cloned into a TA-vector by Mighty TA-cloning Kit (Takara). After LacZ selection, the PCR-amplified inserted products from white colonies were analyzed by sequencing using the *LYRM2* forward primer to determine the deletion size. For the deletion assay in asynchronous cells, the H1299 dA-3 cells (Ogiwara et al., 2011) transfected with indicated siRNA were further transfected with the I-*Sce*I plasmid to induce DSBs and incubated for 2 days. The repair products were amplified from genomic DNA using Ex-Taq (Takara) and cloned into the pCR2.1 vector (Thermo Fisher Scientific). After LacZ selection, the PCR-amplified inserted products from white colonies were analyzed by polyacrylamide gel electrophoresis (PAGE). For clones identified as having deletions, the size was analyzed by sequencing.

### Chromosome translocation assay

Since p53-dependent G1 arrest limits the number of IR-induced dicentric chromosomes appearing in mitosis (Yamauchi et al., 2011), mitotic cells were harvested in the presence of p53 shRNA. The lentiviral vector that encodes p53 shRNA (shp53 pLKO.1 Puro) was a kind gift from Robert A. Weinberg. Before the chromosome experiment, p53 shRNA was lentivirally introduced into the cells, and p53 shRNA-expressing cells were selected in medium containing puromycin (10 µg mL^-1^) for >2 weeks. For the chromosome experiment, the cells were plated at high density (1.0 × 10^6^ cells per 60 mm dish) and contact-inhibited for 3 days to synchronize in the G1 phase. The cells were then irradiated with 2 Gy γ-rays using a Cs-137 γ-ray irradiator (Pony Industries) at a dose rate of 1 Gy m^-1^ and kept at high density for 24 hours to allow DSB repair in G1. The cells were harvested and replated at low density to allow cell cycle progression to mitosis. After addition of Colcemid (0.025 μg mL^-1^ final) at 12 hours after replating, metaphase cells were harvested at 28 hours after replating, washed once with PBS and subjected to hypotonic treatment in 0.075 M KCl for 20 min at room temperature. The cells were fixed with Carnoy’s fixative (methanol:acetic acid = 3:1) for 30 min ∼ overnight at 4°C, resuspended in the fixative, dropped onto slide glasses and dried overnight. The dried slide glasses were washed briefly in PBS, immersed in 4% paraformaldehyde in PBS for 2 min at room temperature, and washed three times in PBS. Peptide nucleic acid probes for centromeres and telomeres (Panagene) were applied onto the cells, and the slide glasses were heated for 3 min at 80°C to denature DNA, followed by hybridization in the dark for 2 hours at room temperature. After hybridization, the slide glasses were washed twice in wash buffer I (70% formamide in TE buffer) and then, washed twice in wash buffer II (TE buffer containing 0.15 M NaCl and 0.05% Tween 20). The washed slide glasses were briefly air dried and counterstained with SlowFade Antifade Reagent containing DAPI (Thermo Fisher Scientific).

### siRNA, nucleotide, plasmid, and transfection

The siRNAs, oligonucleotides and antibodies used in this study are listed in Tables S2-4. The plasmids for RAP80, CtIP, MSH2, MSH3 or RPA2 expression were generated by inserting the PCR-amplified cDNA fragment into the pEGFP-C1 or pcDNA3.1 vector. The plasmids for GFP-HB, GFP-RAD52, GFP-XPG or RNase H1 expression were described previously (Yasuhara et al., 2018). Cells were transfected with the indicated siRNAs using DharmaFECT (GE healthcare), HiPerFect (Qiagen) or RNAiMAX (Thermo Fisher Scientific). The knockdown efficiency was confirmed by western blot and shown in Figure S7. For plasmid transfection, ViaFect (Promega) or Lipofectamine 2000 (Thermo Fisher Scientific) was used according to the manufacturer’s instructions.

### Database analysis

The normalized RNA sequencing data, deletions in genes and gene fusions for each samples provided by TCGA BRCA project were obtained as described previously (Yasuhara et al., 2018) and organized according to the expression levels of the gene of interest. The list of repair genes analyzed in this study is provided in Table S1. The groups were divided as the ratio (high/low) of median expression of the gene was the lowest or not less than 1.5. The data for cell-cycle regulated genes were obtained from Cyclebase3.0 (https://cyclebase.org/) (Santos et al., 2015).

### Statistics

In the box plots, center lines show the medians; box limits indicate the 25th and 75th percentiles as determined by R software; whiskers extend 1.5 times the interquartile range from the 25^th^ and 75th percentiles; all data points are shown. In this study, two-tailed Welch’s *t*-test was used unless otherwise stated. If necessary, *P*-values were corrected according to the Bonferroni method. The significance levels are shown in each panel as follows. * *P* < 0.05, ** *P* < 0.01, *** *P* < 0.001, \*\*\*\**P* < 0.0001, n.s., not significant.

