## Supplemental Information for "RAP80 suppresses the vulnerability of R-loops during DNA double-strand break repair"

### Supplemental figure legends

#### Figure S1. RAP80 suppresses deletion size during G1 DSB repair at transcriptionally active regions.

(A) Sequencing analysis of deletions at the *LYRM2* locus from the experiments shown in Figure 1E. The original sequence is shown in green. (B) A schematic representation of the deletion analysis at a transcriptionally active locus in asynchronous H1299 dA-3 cells. (C) The effect of RAP80 depletion on deletions within transcriptionally active loci was analyzed in asynchronous H1299 dA-3 cells.

#### Figure S2. RNF168 and TRAIP-independent recruitment of RAP80 to DNA damage.

(A, B) The effect of RNF168 (A) or TRAIP (B) depletion on RAP80 recruitment to the laser track was analyzed in G1 U2OS cells (mean ± SEM., *n*=3 for the data points of three biological replicates).

#### Figure S3. RAP80 promotes recruitment of XPG to DNA damage.

(A, B) The effect of MDC1 or RNF168 (A) or BRCA1 or CtIP (B) depletion on DNA-RNA hybrid accumulation at the laser track was analyzed in G1 U2OS cells (mean ± SEM., *n*=3 for the data points of three biological replicates). (C) The results of control experiments (-/+DSB induction, -/+RNase H treatment) for DRIP assay are shown (mean ± SD, *n*=2-5 for data points from two biological replicates). No signal was detected in the isotype IgG pull down (ND). (D) The effect of RAD52 or XPG depletion on DNA-RNA hybrid accumulation at the laser track was analyzed in G1 U2OS cells (mean ± SEM., *n*=3 for data points of three biological replicates). (E, F) The effect of RAP80 depletion on RAD52 (E) or XPG (F) recruitment to the laser track was analyzed in G1 U2OS cells (mean ± SEM., *n*=3 for data points of three biological replicates). (G) The effect of RAD52 or XPG depletion on CtIP recruitment to the laser track was analyzed in G1 U2OS cells (mean ± SEM., *n*=3 for data points of three biological replicates). (H) The effect of exogenous RNase H1 expression on CtIP recruitment to the laser track in RAD52- or XPG-depleted cells was analyzed in G1 U2OS cells (mean ± SEM., *n*=3 for data points of three biological replicates).

**Figure S4. RPA is not recruited at early time points after DNA damage induction in G1 cells.**

(A) The effect of S1 nuclease or RNase A treatment on the IF signals of ssDNA or DNA stem loop was analyzed at 1 min after laser irradiation in U2OS cells. The irradiated lines are indicated with the white arrows. Scale bars, 10 µm. (B, C) The effect of TRi on ssDNA (B) or DNA stem loop (C) accumulation 1 min after laser irradiation was analyzed by IF in U2OS cells (mean ± SEM., *n*=25, 38 (c, DMSO, TRi), *n*=53, 34 (d, DMSO, TRi) for data points of independent cells from three biological replicates). (D) The accumulation of GFP-RPA2 at the laser track was analyzed in G1 U2OS cells. The images at the indicated time points from one representative analysis are shown. White arrowheads, the position of laser irradiation. Scale bars, 10 µm. (E, F) The effect of RAD52 or XPG (E) or CtIP (F) depletion on MSH2 recruitment to the laser track was analyzed in G1 U2OS cells (mean ± SEM., *n*=3 for data points of three biological replicates).

**Figure S5. RAP80, BRCA1, and Polθ are involved in TA-EJ in G1 cells.**

(A) An example of the variation between three biological replicates of the γH2AX foci analysis. One representative result is shown in Figure 6A (mean ± SEM., *n*=3 for the data points of three biological replicates). (B) The results of foci analysis in Figure 6A (30 min time point) were reanalyzed by an automated focus counting method using ImageJ software or blindly counted by a researcher from an independent laboratory (mean with 95% CI, *n*=286, 188, 30, 30, left to right, for data points of independent cells). (C) The effect of CtIP depletion on DSB repair efficiency was analyzed in wild-type or RAP80-depleted cells by measuring the number of γH2AX foci at 30 min after IR in G1 RPE cells. Representative results from three biological replicates are shown (mean with 95% CI, *n*=25 for data points of independent cells). (D, E) The effect of Polθ (D) or LIG1/3 (E) depletion on DSB repair efficiency in BRCA1-depleted cells was analyzed by measuring the number of γH2AX foci 30 min after IR in G1 RPE cells. Representative results from three biological replicates are shown (mean with 95% CI, *n*=25 for data points of independent cells).

**Figure S6. CtIP is involved in TA-EJ.**

(A) The effect of CtIP depletion on DSB repair efficiency in BRCA1-depleted cells was analyzed by measuring the number of γH2AX foci 30 min after IR in G1 RPE cells. Representative results from three biological replicates are shown (mean with 95% CI, *n*=25 for data points of independent cells). (B) The impact of the CtIP mutants on DSB repair efficiency was analyzed as in (A). (C) The effect of CtIP depletion on chromosome translocation frequency was analyzed by counting the number of dicentric chromosomes per cell. Over 2,000 mitotic chromosomes from 50 independent cells were scored per condition in total (mean ± SD, $n$=3 for the data points of three biological replicates). (D) The effect of RAP80 or BRCA1 depletion on DSB repair efficiency in RAD52- or XPG-depleted cells was analyzed as in (A). (E) The effect of Artemis depletion on DSB repair efficiency in RAD52- or XPG-depleted cells was analyzed as in (A). (F) The effect of exogenous RNase H1 expression on DSB repair efficiency in RAD52- or XPG-depleted ΔArtemis cells was analyzed as in (A).

**Figure S7. Confirmation of knockout and knockdown efficiency.**

The knockdown efficiency and knockout were confirmed by western blot. The band for RAP80 or a non-specific protein is indicated by an arrow or asterisk, respectively.

A

RPE Deletion assay in G1
*LYRM2* locus

siControl (*N*=61)

AsiSI site

```
No. GGGAAATCAAGCGCCTGACGCTGTCTCACGCGAT|CGCACCAAACCAAGGAAGCAGCCAGTGCCTC DEL size (bp)
 1 GGGAAATCAAGCGCCTGACGCTGTCTCACGCG-TCGCACCAAACCAAGGAAGCAGCCAGTGCCTC    1
 2 GGGAAATCAAGCGCCTGACGCTGTCTCACGCG-TCGCACCAAACCAAGGAAGCAGCCAGTGCCTC    1
 3 GGGAAATCAAGCGCCTGACGCTGTCTCACGCG-TCGCACCAAACCAAGGAAGCAGCCAGTGCCTC    1
 4 GGGAAATCAAGCGCCTGACGCTGTCTCACGG-ATCGCACCAAACCAAGGAAGCAGCCAGTGCCTC    1
 5 GGGAAATCAAGCGCCTGACGCTGTCTCACG-GATCGCACCAAACCAAGGAAGCAGCCAGTGCCTC    1
 6 GGGAAATCAAGCGCCTGACGCTGTCTCACG-GATCGCACCAAACCAAGGAAGCAGCCAGTGCCTC    1
 7 GGGAAATCAAGCGCCTGACGCTGTCTCACG-GATCGCACCAAACCAAGGAAGCAGCCAGTGCCTC    1
 8 GGGAAATCAAGCGCCTGACGCTGTCTCACG-GATCGCACCAAACCAAGGAAGCAGCCAGTGCCTC    1
 9 GGGAAATCAAGCGCCTGACGCTGTCTCACG-GATCGCACCAAACCAAGGAAGCAGCCAGTGCCTC    1
10 GGGAAATCAAGCGCCTGACGCTGTCTCACG-GATCGCACCAAACCAAGGAAGCAGCCAGTGCCTC    1
11 GGGAAATCAAGCGCCTGACGCTGTCTCACG-GATCGCACCAAACCAAGGAAGCAGCCAGTGCCTC    1
12 GGGAAATCAAGCGCCTGACGCTGTCTCACG-GATCGCACCAAACCAAGGAAGCAGCCAGTGCCTC    1
13 GGGAAATCAAGCGCCTGACCCTGTCTCAC-CGATCGCACCACACCAGGACCCGGCCGGTGCCTC    1
14 GGGAAATCAAGCGCCTGACGCTGTCTCACGC--TCGCACCAAACCAAGGAAGCAGCCAGTGCCTC    2
15 GGGAAATCAAGCGCCTGACGCTGTCTCA--CGATCGCACCAAACCAAGGAAGCAGCCAGTGCCTC    2
16 GGGAAATCAAGCGCCTGACGCTGTCTCA--CGATCGCACCAAACCAAGGAAGCAGCCAGTGCCTC    2
17 GGGAAATCAAGCGCCTGACGCTGTCTCA--CGATCGCACCAAACCAAGGAAGCAGCCAGTGCCTC    2
18 GGGAAATCAAGCGCCTGACGCTGTCTCACGCGAT---ACCAAACCAAGGAAGCAGCCAGTGCCTC    3
19 GGGAAATCAAGCGCCTGACGCTGTCTCACGCGAT--AT-CAAACCAAAGAAGCGACCCGTGCCTC    3
20 GGGAAATCAAGCGCCTGACGCTGTCTCACGCGAT---ACCAAACCAAGGAAGCAGCCAGTGCCTC    3
21 GGGAAATCAAGCGCCTGACGCTGTCTCACGCGA---CACCAAACCAAGGAAGCAGCCAGTGCCTC    3
22 GGGAAATCAAGCGCCTGACGCTGTCTCACGCGA---CACCAAACCAAGGAAGCAGCCAGTGCCTC    3
23 GGGAAATCAAGCGCCTGACGCTGTCTCACGCGA---CACCAAACCAAGGAAGCAGCCAGTGCCTC    3
24 GGGAAATCAAGCGCCTGACGCTGTCTCACGCGA---CACCAAACCAAGGAAGCAGCCAGTGCCTC    3
25 GGGAAATCAAGCGCCTGACGCTGTCTCACGCGA---CACCAAACCAAGGAAGCAGCCAGTGCCTC    3
26 GGGAAATCAAGCGCCTGACGCTGTCTCACG---TCGCACCAAACCAAGGAAGCAGCCAGTGCCTC    3
27 GGGAAATCAAGCGCCTGACGCTGTCTCACG---TCGCACCAAACCAAGGAAGCAGCCAGTGCCTC    3
28 GGGAAATCAAGCGCCTGACGCTGTCTCACG---TCGCACCAAACCAAGGAAGCAGCCAGTGCCTC    3
29 GGGAAATCAAGCGCCTGACGCTGTCTCACG---TCGCACCAAACCAAGGAAGCAGCCAGTGCCTC    3
30 GGGAAATCAAGCGCCTGACGCTGTCTCAC---ATCGCACCAAACCAAGGAAGCAGCCAGTGCCTC    3
31 GGGAAATCAAGCGCCTGACGCTGTCTCAC---ATCGCACCAAACCAAGGAAGCAGCCAGTGCCTC    3
32 GGGAAATCAAGCGCCTGACGCTGTCTCA--A-ATCGCACCAAACCAAGGAAGCAGCCAGTGCCTC    3
33 GGGAAATCAAGCGCCTGACGCTGTCTCA--A-ATCGCACCAAACCAAGGAAGCAGCCAGTGCCTC    3
34 GGGAAATCAAGCGCCTGACGCTGTCTCA--T-ATCGCACCAAACCAAGGAAGCAGCCAGTGCCTC    3
35 GGGAAATCAAGCGCCTGACGCTGTCTCA--T-ATCGCACCAAACCAAGGAAGCAGCCAGTGCCTC    3
36 GGGAAATCAAGCGCCTGACGCTGTCTCAC----TCGCACCAAAACAAGGAAGCAGCCTGTGCCTC    4
37 GGGAAATCAAGCGCCTGACGCTGTCTCA----ATCGCACCAAACCAAGGAAGCAGCCAGTGCCTC    4
38 GGGAAATCAAGCGCCTGACGCTGTCTCA----ATCGCACCAAACCAAGGAAGCAGCCAGTGCCTC    4
39 GGGAAATCAAGCGCCTGACGCTGTCTCA----ATCGCACCAAACCAAGGAAGCAGCCAGTGCCTC    4
40 GGGAAATCAAGCGCCTGACGCTGTCTCA----ATCGCACCAAACCAAGGAAGCAGCCAGTGCCTC    4
41 GGGAAATCAAGCGCCTGACGCTGTCTCA----ATCGCACCAAACCAAGGAAGCAGCCAGTGCCTC    4
42 GGGAAATCAAGCGCCTGACGCTGTCTCA-----TCGCACCAAACCAAGGAAGCAGCCAGTGCCTC    5
43 GGGAAATCAAGCGCCTGACGCTGTCTCACGCG---T---CAAACCAAGGAAGCAGCCAGTGCCTC    6
44 GGGAAATCAAGCGCCTGACGCTGTCTCACGCG---T---CAAACCAAGGAAGCAGCCAGTGCCTC    6
45 GGGAAATCAAGCGCCTGACGCTGTCTCACGC------ACCAAACCAAGGAAGCAGCCAGTGCCTC    6
46 GGGAAATCAAGCGCCTGACGCTGTCTCACGC------ACCAAACCAAGGAAGCAGCCAGTGCCTC    6
47 GGGAAATCAAGCGCCTGATCCTGTCTCACGC------ACCAAACCAAGGAAGCCTCCCGTGCCGC    6
48 GGGAAATCAAGCGCCTGACGCTGTCTCACGC------ACCAAACCAAGGAAGCAGCCAGTGCCTC    6
49 GGGAAATCAAGCGCCTGACGCTGTCTCACGC------ACCAAACCAAGGAAGCAGCCAGTGCCTC    6
50 GGGAAATCAAGCGCCTGACGCTGTCTCACGC------ACCAAACCAAGGAAGCAGCCAGTGCCTC    6
51 GGGAAATCAAGCGCCTGACGCTGTCTCACGC------ACCAAACCAAGGAAGCAGCCAGTGCCTC    6
52 GGGAAATCAAGCGCCTGACGCTGTCTCACGCG---------AACCAAGGAAGCAGCCAGTGCCTC    9
53 GGGAAATCAAGCGCCTGACGCTGTCTCACGCG---------AACCAAGGAAGCAGCCAGTGCCTC    9
54 GGGAAATCAAGCGCCTGACGCTGTCTCACGCGAT------------AGGAAGCAGCCAGTGCCTC   12
55 GGGAAATCAAGCGCCTGACG-------------TCGCACCAAACCAAGGAAGCAGCCAGTGCCTC   13
56 GGGAAATCAAGCGCCTGACG-------------TCGCACCAAACCAAGGAAGCAGCCAGTGCCTC   13
57 GGGAAATCAAGCGCCTGA--------------ATCGCACCAAACCAAGGAAGCAGCCAGTGCCTC   14
58 GGGAAATCAAGCGCCTGACGCTGTCTCACGCGAT---------------AAGCAGCCAGTGCCTC   15
59 GGGAAATCAAGCGCCTGACGCTGTCTCACGCGAT---------------AAGCAGCCAGTGCCTC   15
60 GGGAAATCAAGCGCCTGACGCTGTCTCACGCGAT---------------AAGCAGCCAGTGCCTC   15
61 GGGAAATCAAGCGCCTGA---------------TCGCACCAAACCAAGGAAGCAGCCAGTGCCTC   15
62 GGGAAATCAAGCGCCTGA---------------TCGCACCAAACCAAGGAAGCAGCCAGTGCCTC   15
63 GGGAAATCAAGCGCCTGACGCTGTCTCACGCGAT------------------CAGCCAGTGCCTC   18
64 GGGAAATCAAGCGCCTGACGCTGTCT--------------------------CAGCCAGTGCCTC   26
65 --(43 bp del) ---------------------GCACCAAACCAAGGAAGCAGCCAGTGCCTC   43
```

siRAP80 (*N*=56)

AsiSI site

```
No. GGGAAATCAAGCGCCTGACGCTGTCTCACGCGAT|CGCACCAAACCAAGGAAGCAGCCAGTGCCTC DEL size (bp)
 1 GGGAAATCAAGCGCCTGACGCTGTCTCACGCGAT-GCACCAAACCAAGGAAGCAGCCAGTGCCTC    1
 2 GGGAAATCAAGCGCCTGACGCTGTCTCACGCG-TCGCACCAAACCAAGGAAGCAGCCAGTGCCTC    1
 3 GGGAAATCAAGCGCCTGACGCTGTCTCACGCG-TCGCACCAAACCAAGGAAGCAGCCAGTGCCTC    1
 4 GGGAAATCAAGCGCCTGACGCTGTCTCACGCG-TCGCACCAAACCAAGGAAGCAGCCAGTGCCTC    1
 5 GGGAAATCAAGCGCCTGACGCTGTCTCACGC-ATCGCACCAAACCAAGGAAGCAGCCAGTGCCTC    1
 6 GGGAAATCAAGCGCCTGACGCTGTCTCACGG-ATCGCACCAAACCAAGGAAGCAGCCAGTGCCTC    1
 7 GGGAAATCAAGCGCCTGACGCTGTCTCACGG-ATCGCACCAAACCAAGGAAGCAGCCAGTGCCTC    1
 8 GGGAAATCAAGCGCCTGACGCTGTCTCACGG-ATCGCACCAAACCAAGGAAGCAGCCAGTGCCTC    1
 9 GGGAAATCAAGCGCCTGACGCTGTCTCACGC-ATCGCACCAAACCAAGGAAGCAGCCAGTGCCTC    1
10 GGGAAATCAAGCGCCTGACGCTGTCTCACGC-ATCGCACCAAACCAAGGAAGCAGCCAGTGCCTC    1
11 GGGAAATCAAGCGCCTGACGCTGTCTCACG-GATCGCACCAAACCAAGGAAGCAGCCAGTGCCTC    1
12 GGGAAATCAAGCGCCTGACGCTGTCTCACG-GATCGCACCAAACCAAGGAAGCAGCCAGTGCCTC    1
13 GGGAAATCAAGCGCCTGACGCTGTCTCACG-GATCGCACCAAACCAAGGAAGCAGCCAGTGCCTC    1
14 GGGAAATCAAGCGCCTGACGCTGTCTCACG-GATCGCACCAAACCAAGGAAGCAGCCAGTGCCTC    1
15 GGGAAATCAAGCGCCTGACGCTGTCTCACG--ATCGCACCAAACCAAGGAAGCAGCCAGTGCCTC    2
16 GGGAAATCAAGCGCCTGACGCTGTCTCA--CGATCGCACCAAACCAAGGAAGCAGCCAGTGCCTC    2
17 GGGAAATCAAGCGCCTGACGCTGTCTCACGCGAT---ACCAAACCAAGGAAGCAGCCAGTGCCTC    3
18 GGGAAATCAAGCGCCTGACGCTGTCTCACG-A--CGCACCAAACCAAGGAAGCAGCCAGTGCCTC    3
19 GGGAAATCAAGCGCCTGACGCTGTCTCACG-A--CGCACCAAACCAAGGAAGCAGCCAGTGCCTC    3
20 GGGAAATCAAGCGCCTGACGCTGTCTCACGC---CGCACCAAACCAAGGAAGCAGCCAGTGCCTC    3
21 GGGAAATCAAGCGCCTGACGCTGTCTCACG----CGCACCAAACCAAGGAAGCAGCCAGTGCCTC    4
22 GGGAAATCAAGCGCCTGACGCTGTCTCA----ATCGCACCAAACCAAGGAAGCAGCCAGTGCCTC    4
23 GGGAAATCAAGCGCCTGACGCTGTCTCA----ATCGCACCAAACCAAGGAAGCAGCCAGTGCCTC    4
24 GGGAAATCAAGCGCCTGACGCTGTCTCA----ATCGCACCAAACCAAGGAAGCAGCCAGTGCCTC    4
25 GGGAAATCAAGCGCCTGACGCTGTCTCA----ATCGCACCAAACCAAGGAAGCAGCCAGTGCCTC    4
26 GGGAAATCAAGCGCCTGACGCTGTCTCA----ATCGCACCAAACCAAGGAAGCAGCCAGTGCCTC    4
27 GGGAAATCAAGCGCCTGACGCTGTCTCA----ATCGCACCAAACCAAGGAAGCAGCCAGTGCCTC    4
28 GGGAAATCAAGCGCCTGACGCTGTCTCA----ATCGCACCAAACCAAGGAAGCAGCCAGTGCCTC    4
29 GGGAAATCAAGCGCCTGACGCTGTCTCA----ATCGCACCAAACCAAGGAAGCAGCCAGTGCCTC    4
30 GGGAAATCAAGCGCCTGACGCTGTCTCA----ATCGCACCAAACCAAGGAAGCAGCCAGTGCCTC    4
31 GGGAAATCAAGCGCCTGACGCTGTCTCA----ATCGCACCAAACCAAGGAAGCAGCCAGTGCCTC    4
32 GGGAAATCAAGCGCCTGACGCTGTCTCACGCGAT-----CAAACCAAGGAAGCAGCCAGTGCCTC    5
33 GGGAAATCAAGCGCCTGACGCTGTCTCAC-----CGCACCAAACCAAGGAAGCAGCCAGTGCCTC    5
34 GGGAAATCAAGCGCCTGACGCTGTCTC-----ATCGCACCAAACCAAGGAAGCAGCCAGTGCCTC    5
35 GGGAAATCAAGCGCCTGACGCTGTCTCA------CGCACCAAACCAAGGAAGCAGCCAGTGCCTC    6
36 GGGAAATCAAGCGCCTGACGCTGTCTC-------CGCACCAAACCAAGGAAGCAGCCAGTGCCTC    7
37 GGGAAATCAAGCGCCTGACGCTGTCTC-------CGCACCAAACCAAGGAAGCAGCCAGTGCCTC    7
38 GGGAAATCAAGCGCCTGACGCTGT--------ATCGCACCAAACCAAGGAAGCAGCCAGTGCCTC    8
39 GGGAAATCAAGCGCCTGACGCTGTCTCAC----------CAAACCAAGGAAGCAGCCAGTGCCTC   10
40 GGGAAATCAAGCGCCTGACGCTGTCT----------CACCAAACCAAGGAAGCAGCCAGTGCCTC   10
41 GGGAAATCAAGCGCCTGACGCTGTCT----------CACCAAACCAAGGAAGCAGCCAGTGCCTC   10
42 GGGAAATCAAGCGCCTGACGCTGTC----------GCACCAAACCAAGGAAGCAGCCAGTGCCTC   10
43 GGGAAATCAAGCGCCTGACGCTGTC----------GCACCAAACCAAGGAAGCAGCCAGTGCCTC   10
44 GGGAAATCAAGCGCCTGACGCT-CGG-----------ACCAAACCAAGGAAGCAGCCAGTGCCTC   12
45 GGGAAATCAAGCGCCTGAT---------------CGCACCAAACCAAGGAAGCAGCCAGTGCCTC   15
46 GGGAAATCAAGCGCCTGAT---------------CGCACCAAACCAAGGAAGCAGCCAGTGCCTC   15
47 GGGAAATCAAGCGCCTGA---------------TCGCACCAAACCAAGGAAGCAGCCAGTGCCTC   15
48 GGGAAATCAAGCGCCTGACGCTGTCT------------------CAAGGAAGCAGCCAGTGCCTC   18
49 GGGAAATCAAGCGCCTGACGCTGTCTCACGC-----------------------GCCAGTGCCTC   23
50 GGGAAATCAA-----------------------TCGCACCAAACCAAGGAAGCAGCCAGTGCCTC   23
51 GGGAAATCAAGCGCCTGACGCTGTCTCACG-A------------------------CAGTGCCTC   25
52 GGGAAATCAAGCGCCTGACGCTGTCTCACG-A------------------------CAGTGCCTC   25
53 GGGAAATCAA-------------------------GCACCAAACCAAGGAAGCAGCCAGTGCCTC   25
54 GGGAAATCAAGCGCC--------------------------AACCAAGGAAGCAGCCAGTGCCTC   26
55 GGGAAATCAAGCGCC--------------------------------GGAAGCAGCCAGTGCCTC   32
56 GGGAAATCAAGCGCCTGACGCTGTCTCACGA-------------(50 bp del)----------   50
```

B

H1299 Deletion assay
(asynchronous)

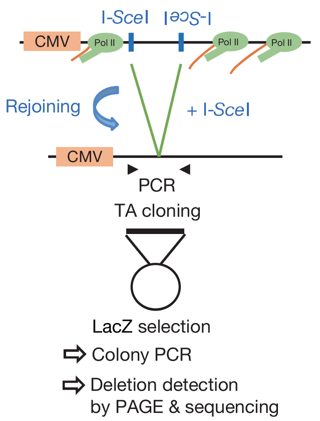


C

H1299 Deletion assay
(asynchronous)

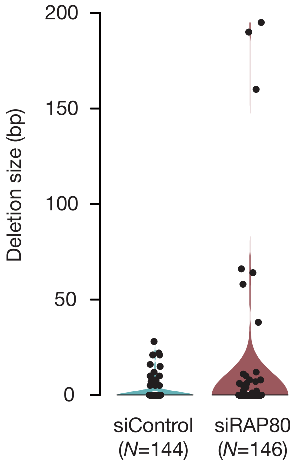

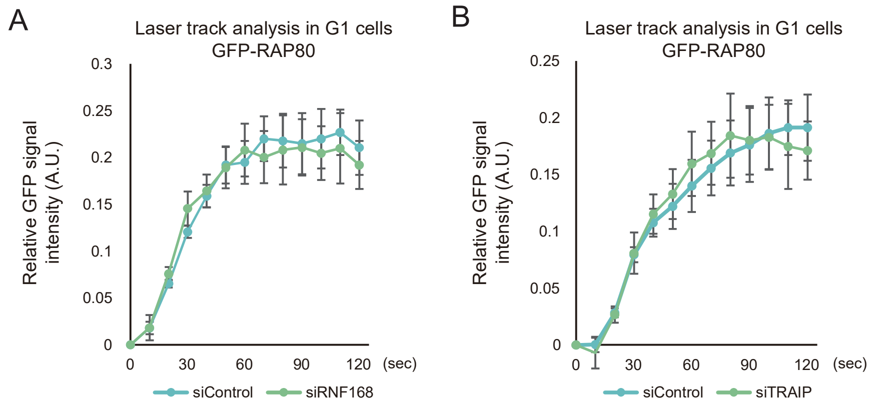
A
Laser track analysis in G1 cells
GFP-RAP80
Relative GFP signal intensity (A.U.)
0
0.05
0.1
0.15
0.2
0.25
0.3
0
30
60
90
120
(sec)
siControl
siRNF168
B
Laser track analysis in G1 cells
GFP-RAP80
Relative GFP signal intensity (A.U.)
0
0.05
0.1
0.15
0.2
0.25
0
30
60
90
120
(sec)
siControl
siTRAIP

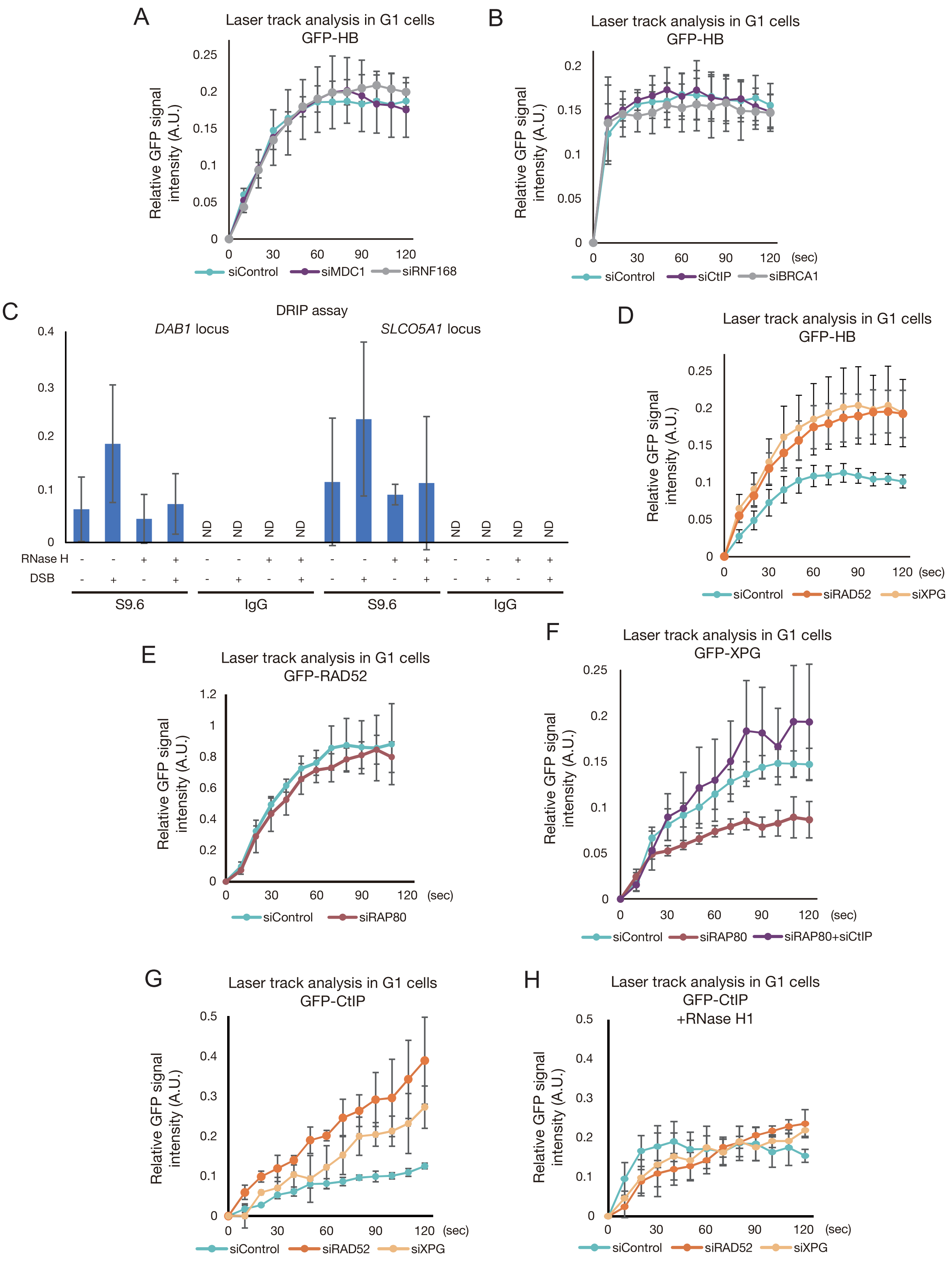
A
Laser track analysis in G1 cells
GFP-HB
Relative GFP signal intensity (A.U.)
0 0.05 0.1 0.15 0.2 0.25
0 30 60 90 120
siControl siMDC1 siRNF168
B
Laser track analysis in G1 cells
GFP-HB
Relative GFP signal intensity (A.U.)
0 0.05 0.1 0.15 0.2
0 30 60 90 120 (sec)
siControl siCtIP siBRCA1
C
DRIP assay
DAB1 locus
SLCO5A1 locus
0 0.1 0.2 0.3 0.4
ND ND ND ND ND ND ND ND
RNase H - - + + - - + + - - + + - - + +
DSB - + - + - + - + - + - + - + - +
S9.6 IgG S9.6 IgG
D
Laser track analysis in G1 cells
GFP-HB
Relative GFP signal intensity (A.U.)
0 0.05 0.1 0.15 0.2 0.25
0 30 60 90 120 (sec)
siControl siRAD52 siXPG
E
Laser track analysis in G1 cells
GFP-RAD52
Relative GFP signal intensity (A.U.)
0 0.2 0.4 0.6 0.8 1 1.2
0 30 60 90 120 (sec)
siControl siRAP80
F
Laser track analysis in G1 cells
GFP-XPG
Relative GFP signal intensity (A.U.)
0 0.05 0.1 0.15 0.2 0.25
0 30 60 90 120 (sec)
siControl siRAP80 siRAP80+siCtIP
G
Laser track analysis in G1 cells
GFP-CtIP
Relative GFP signal intensity (A.U.)
0 0.1 0.2 0.3 0.4 0.5
0 30 60 90 120 (sec)
siControl siRAD52 siXPG
H
Laser track analysis in G1 cells
GFP-CtIP
+RNase H1
Relative GFP signal intensity (A.U.)
0 0.1 0.2 0.3 0.4 0.5
0 30 60 90 120 (sec)
siControl siRAD52 siXPG

Yasuhara_Fig. S4

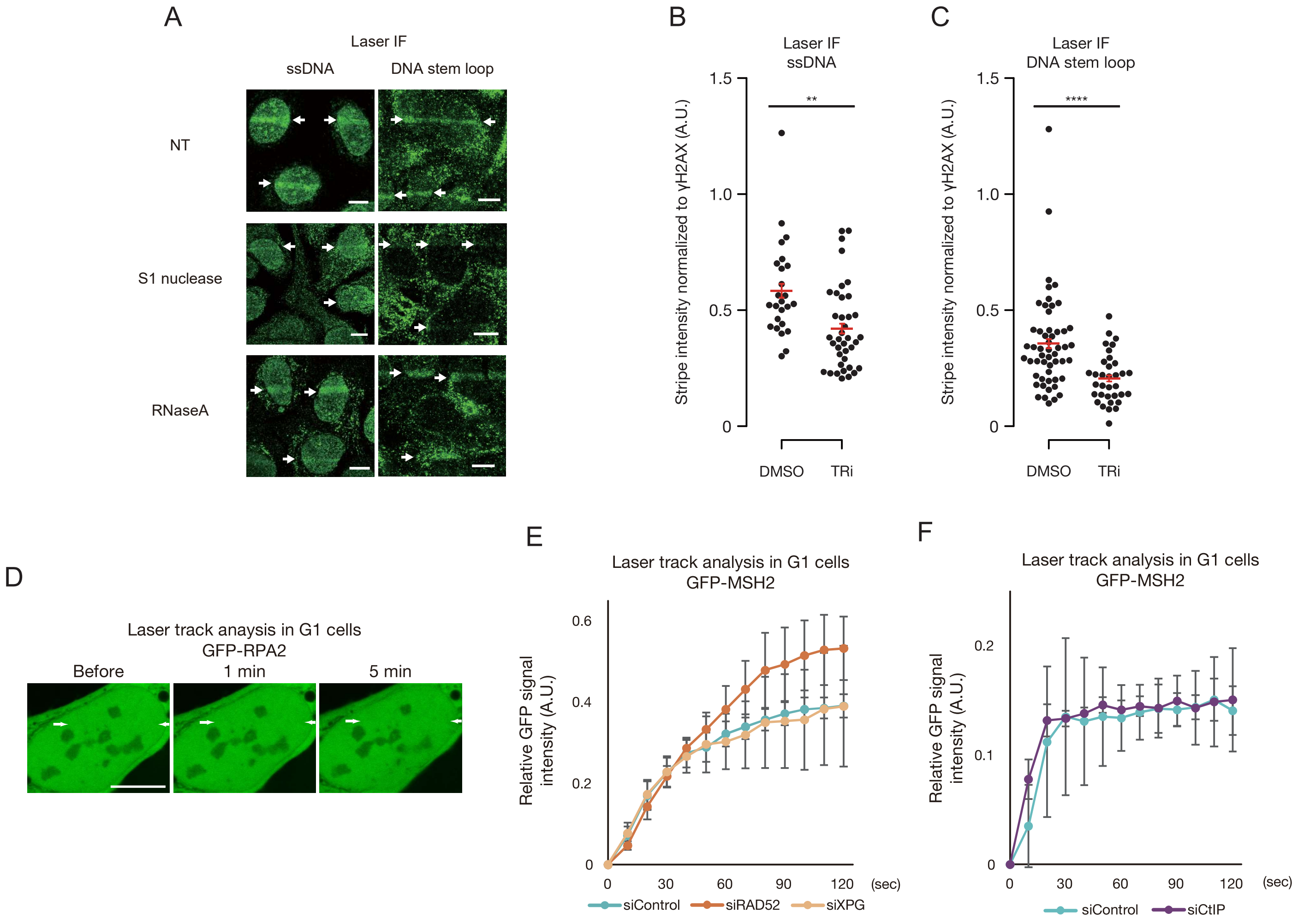

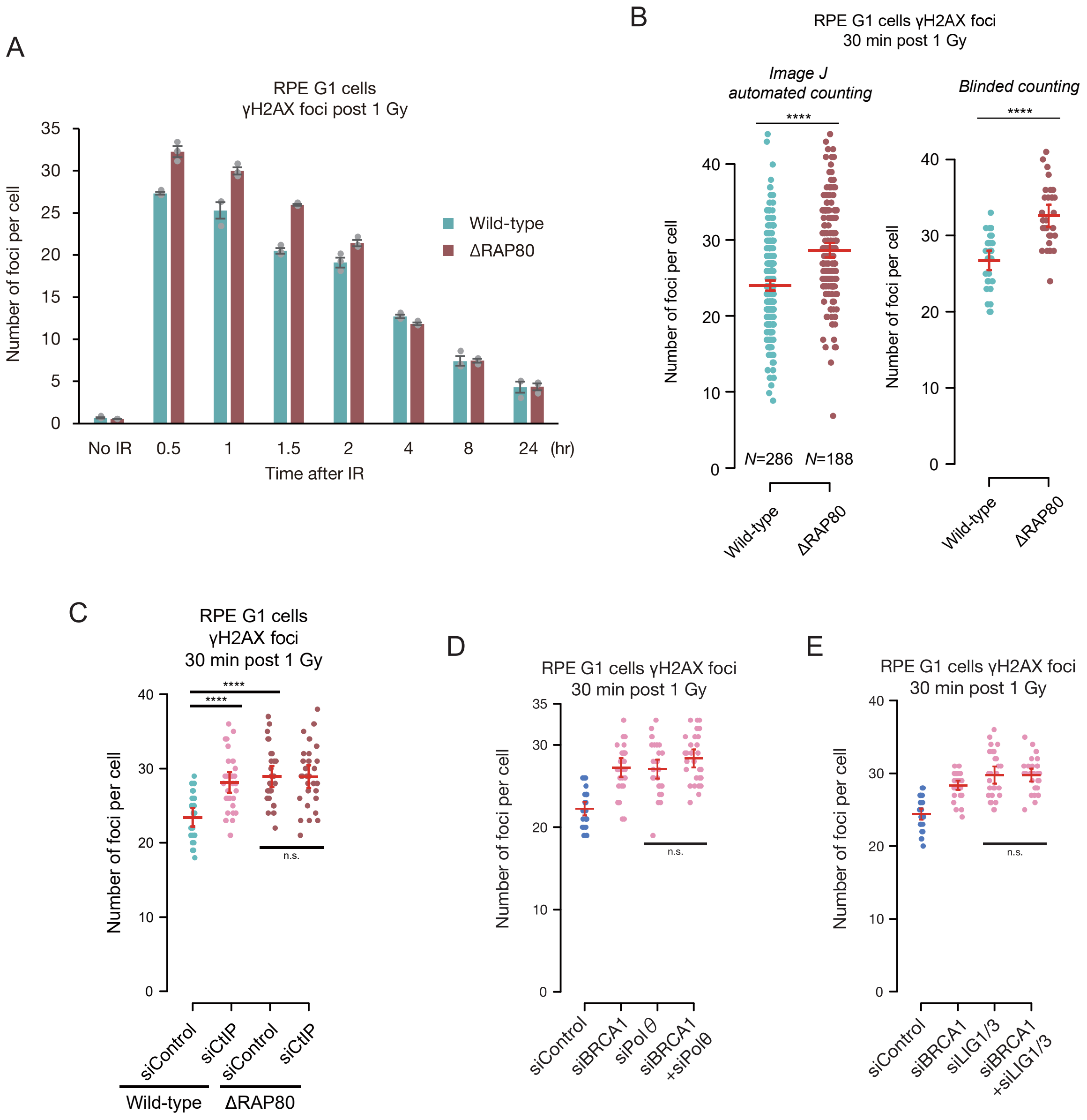
A
RPE G1 cells
γH2AX foci post 1 Gy
Number of foci per cell
35
30
25
20
15
10
5
0
Wild-type
ΔRAP80
No IR
0.5
1
1.5
2
4
8
24
(hr)
Time after IR
B
RPE G1 cells γH2AX foci
30 min post 1 Gy
Image J
automated counting
Blinded counting
****
****
Number of foci per cell
40
30
20
10
0
N=286
N=188
Wild-type
ΔRAP80
C
RPE G1 cells
γH2AX foci
30 min post 1 Gy
****
****
n.s.
siControl
siCtIP
Wild-type
ΔRAP80
D
RPE G1 cells γH2AX foci
30 min post 1 Gy
siBRCA1
siPol θ
siBRCA1
+siPolθ
E
siLIG1/3
+siLIG1/3

Yasuhara_Fig. S6

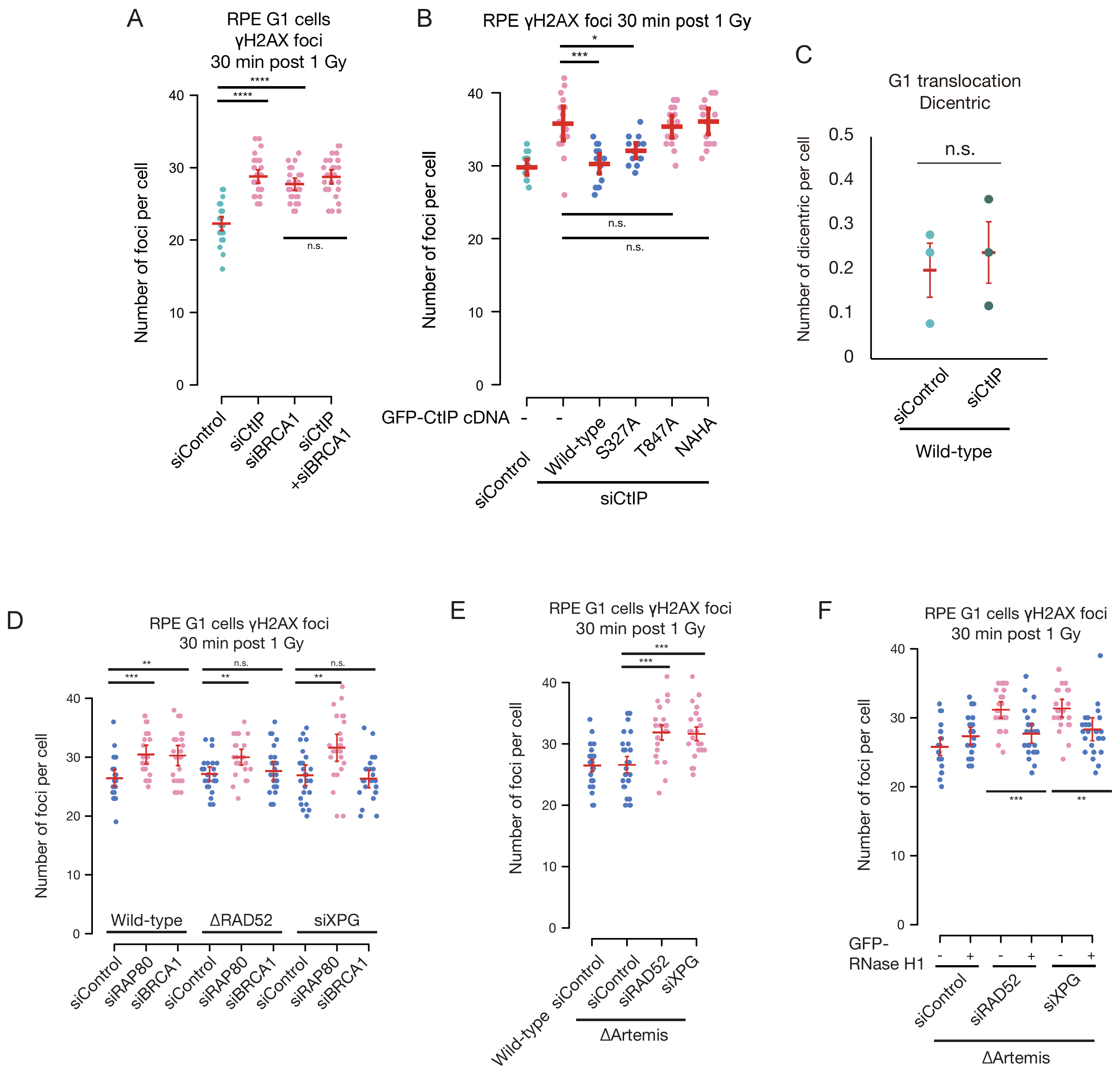

Yasuhara_Fig. S7

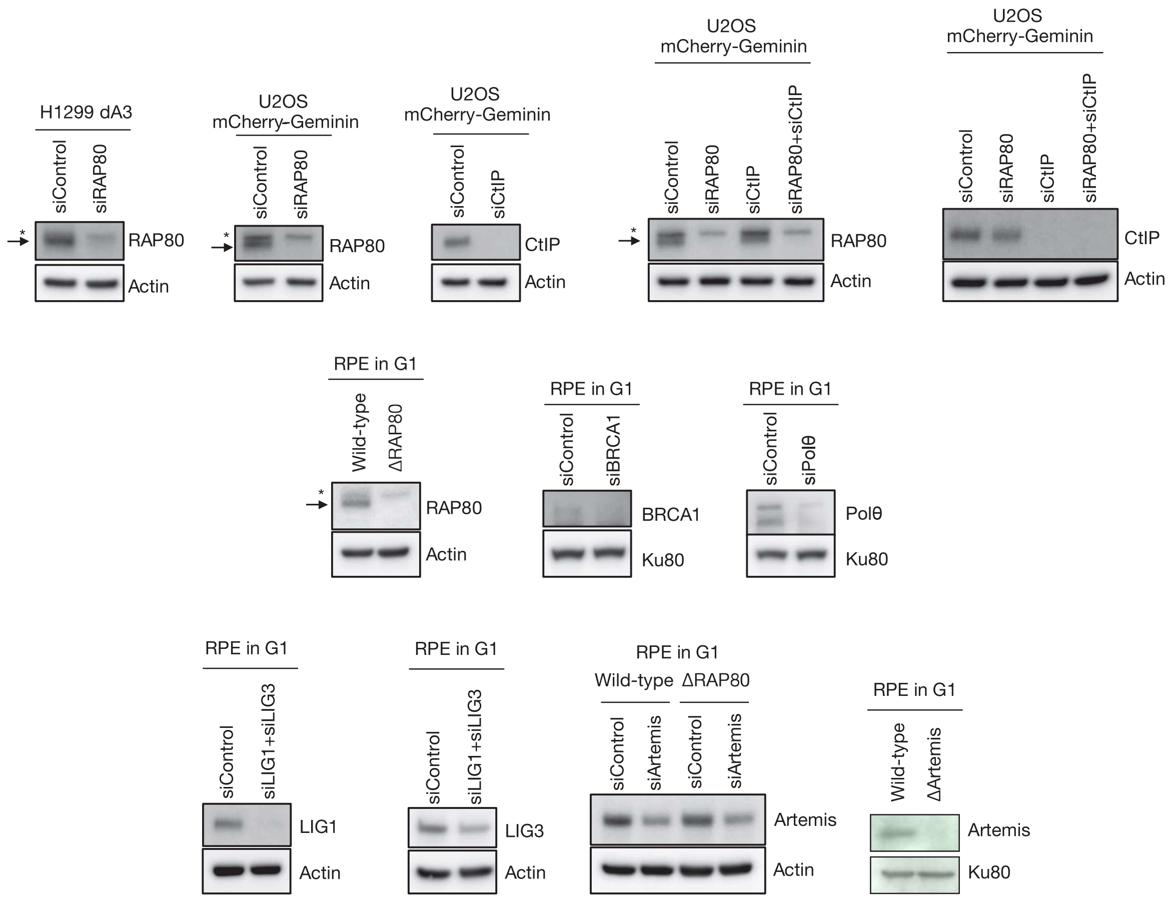

**Table S1. List of repair genes tested in TCGA database analysis.**

| | | |
|---|---|---|
| ATM | FANCE | RAD50 |
| ATR | FANCF | RAD51 |
| ATRIP | FANCG | RAD51AP1 |
| ATRX | FANCI | RAD51B |
| BACH1 | FANCL | RAD51C |
| BARD1 | FANCM | RAD51D |
| BLM | GEN1 | RAD52 |
| BRCA1 | H2AFX | RAD54B |
| BRCA2 | HELB | RAD54L |
| BRCC3 | KEAP1 | RBBP8 |
| C9orf142 | LIG1 | RIF1 |
| CHEK1 | LIG3 | RNF168 |
| CHEK2 | LIG4 | RNF169 |
| DCLRE1C | MAD2L2 | RNF8 |
| DNA2 | MDC1 | SETD2 |
| DYNLL1 | MRE11 | SLFN11 |
| ERCC1 | MUS81 | SMARCA1 |
| ERCC2 | NBN | SMARCAD1 |
| ERCC3 | NHEJ1 | TOPBP1 |
| ERCC4 | PALB2 | TP53BP1 |
| ERCC5 | PAXIP1 | TRAIP |
| ERCC6 | PBRM1 | UCHL5 |
| ERCC8 | POLQ | UHRF1 |
| EXD2 | PRKDC | UIMC1 |
| EXO1 | PRPF19 | XRCC1 |
| FAM175A | RAD1 | XRCC2 |
| FANCA | RAD18 | XRCC3 |
| FANCB | RAD21 | XRCC4 |
| FANCC | RAD23A | XRCC5 |
| FANCD2 | RAD23B | XRCC6 |

**Table S2. List of siRNAs used in this study.**

| Target | Supplier | Sequence (sense 5'-3') or ID |
|---|---|---|
| Control | Dharmacon | D-001210-05-20 |
| Control | SIGMA | CGUACGCGGAAUACUUCGA |
| RAP80 | Dharmacon | M-006995-03-0005 |
| RAP80 | SIGMA | GAAGGAUGUGGAAACUACCUU |
| CtIP | Dharmacon | L-011376-00-0005 |
| CtIP | SIGMA | GCUAAAACAGGAACGAAUC |
| RAD52 | Dharmacon | M-011760-01-0005 |
| RAD52 | SIGMA | GGAAAUAUGAUCCAUCUUA |
| XPG | SIGMA | GGACUUAGCGUCCAGUGAC |
| RNF168 | Dharmacon | L-007152-00-0005 |
| TRAIP | SIGMA | GCAGCAGGAUGAGACCAAA |
| MDC1 | SIGMA | ACAGUUGUCCCCACAGCCC |
| BRCA1 | Dharmacon | M-003461-02-0005 |
| BRCA1 | SIGMA | UCACAGUGUCCUUUAUGUA |
| POLQ | Dharmacon | M-015180-01-0005 |
| LIG1 | Dharmacon | M-009227-02-0005 |
| LIG3 | Dharmacon | M-004254-00-0005 |
| Artemis | Dharmacon | M-004269-02-0005 |

**Table S3. List of oligonucleotides used in this study.**

| Experiment | Target gene | Oligo name | Sequence |
|---|---|---|---|
| Deletion assay | LYRM2 | LYRM2_F | AGAAAGCATCACGAGGTTCATC |
| | | LYRM2_R | GCGACGCTAACGTTAAAGCA |
| | (H1299) | BP-F1 | GTACGGTGGGAGGTCTATATAAG |
| | | maxGFP-R2 | TTCATCTTGTTGGTCATGCGG |
| DRIP-qPCR | DAB1 | DAB1-5_F | CTCCAGGGCATCCTTAGTGT |
| | | DAB1-5_R | GGGCCAGGTGTGTACTTAGG |
| | SLCO5A1 | SLCO5A1_F | TTCACAGCACTCTCCATTCC |
| | | SLCO5A1_R | TCTTTCCCACCAAGTCTTCA |
| ChIP | LYRM2 | LYRM2-ChIP-F | TCCAAAGCAGCTTACCTGGG |
| | | LYRM2-ChIP-R | CATCGGGCCAATCTCAGAGG |
| | ANP32A | ANP32A-ChIP-F | CAAAGGCTACGTCCCGGTG |
| | | ANP32A-ChIP-R | TATTAAGCAGGCGCCGAATG |
| | SLC32A1 | SLC32A1-ChIP-F | GAACTAGCACCCTAGACGCC |
| | | SLC32A1-ChIP-R | CTCTCAGTAGCCTGGATGCG |
| | Non-gene | NGR-ChIP-F | GGACTGTGTGCCACCCTTAAT |
| | region | NGR-ChIP-R | AAGGAATCAGCTTCACCGGA |
| Cloning | RAP80 | RAP80-F_XhoI | AGATCTCGAGCTATGCCACGGAGAAAGAAA |
| | | RAP80-R_BamHI | CGGTGGATCCTCAGAATTTTCTCCTTCTTCC |
| | | RAP80-UIMD-F | GAAAAATCGCACAGATGTGCCGGCCTTCTGATGC |
| | | RAP80-UIMD-R | GCATCAGAAGGCCGGCACATCTGTGCGATTTTTC |
| | MSH2 | MSH2-F | GCGCATTTTCTTCAACCAG |
| | | MSH2-R | CTGGGATTTTTCACGTAGTAAC |
| | MSH3 | MSH3-F | TCCTTGCCCTGCCATGTCTC |
| | | MSH3-R | GCTCACATGTCACACAAAGATAGG |
| | CtIP | CtIP-F_KpnI | GACGGTACCATGAACATCTCGGGAAGCAGCTGTGG |
| | | CtIP-R_BamHI | GGTGGATCCCTATATGTCTTCTGCTCCTTGCCTTTTGG |
| | CtIP T847A | CtIP-T847A-F | CATTCCACCCAACGCACCAGAGAATTTTTG |
| | | CtIP-T847A-R | CAAAAATTCTCTGGTGCGTTGGGTGGAATG |
| | CtIP NAHA | CtIP-NAHA-F | CCACTGTCTGGAAGGAGCCGCCAAGAAACAGCCTTTTGAGGAATC |
| | | CtIP-NAHA-R | GGCGGCTCCTTCCAGACAGTGGTAGAGCTCATCACCAAGG |
| | RPA2 | RPA2-F | TATGCTCGAGCTATGTGGAACAGTGGATTCGA |
| | | RPA2-R | TCGAGGATCCTGCAGAGCTGGAGACAACAG |

### Table S3 (Continued)

| Experiment | Target gene | Oligo name | Sequence |
|---|---|---|---|
| CRISPR | Artemis | DCLRE1C-ex6-n_AS-F | accgATCTGAAGTCTCCTGTGTAC |
| | | DCLRE1C-ex6-n_AS-R | aaacGTACACAGGAGACTTCAGAT |
| | | DCLRE1C-ex6-n_S-F | accgAATGGAGCTTCTGCACTCCG |
| | | DCLRE1C-ex6-n_S-R | aaacCGGAGTGCAGAAGCTCCATT |
| | | DCLRE1C_ex6_conf-F | TGTAATGGATATGTTTGCAGGAAGC |
| | | DCLRE1C_ex6_conf-R | CCTATACGAGGCCCAGTACC |

**Table S4. List of antibodies used in this study.**

| Target | Cat No. | Company | Application | Dilution |
|---|---|---|---|---|
| Artemis (D7O8V) | 13381 | Cell Signaling | WB | 1:500 |
| BRCA1 | sc-642 | Santa Cruz Biotechnology | WB | 1:100 |
| BrdU (B44) | 347580 | BD Biosciences | IF | 1:100 |
| CtIP (D76F7) | 9201 | Cell Signaling | WB | 1:1,000 |
| DNA stem loop (DNA-1) | Ab00415-1.1 | Absolute Antibody | IF | 1:200 |
| GFP | ab290 | Abcam | ChIP | 1:200 |
| Ku80 (111) | MA5-12933 | Thermo Fisher Scientific | WB | 1:1,000 |
| Ku80 (C48E7) | 2180 | Cell Signaling | WB | 1:2,000 |
| LIG1 | ab177946 | Abcam | WB | 1:1,000 |
| LIG3 | ab185815 | Abcam | WB | 1:1,000 |
| Polθ (1C11) | H00010721-M09 | Novus Biologicals | WB | 1:1,000 |
| RAP80 (D1T6Q) | 14466 | Cell Signaling | IF, WB | IF, 1:100; WB, 1:1,000 |
| RAP80 | A300-763A | Bethyl | IP | 1 µg |
| RNA polymerase II subunit B1 (phospho CTD Ser-2) (3E10) | 04-1571 | Millipore | IP | 1 µg |
| DNA-RNA hybrid (S9.6) | Ab01137-2.0 | Absolute Antibody | IP | 1-10 µg |
| DNA-RNA hybrid (S9.6) | MABE1095 | Millipore | IP | 1 µg |
| ssDNA (TNT-3) | MAB3868 | Millipore | IF | 1:200 |
| ssDNA (TNT-3) | NBP2-29849 | Novus Biologicals | IF | 1:200 |
| β-Actin (8H10D10) | 3700 | Cell Signaling | WB | 1:5,000 |
| γH2AX | A300-081A | Bethyl | IF | 1:2,000 |
| γH2AX (JBW301) | 05-636 | Millipore | IF | 1:1,000 |
